# Interrogation of genome-wide, experimentally dissected gene regulatory networks reveals mechanisms underlying dynamic cellular state control

**DOI:** 10.1101/2021.06.28.449297

**Authors:** Xiangtian Tan, Jeremy Worley, Mikko Turunen, Kelly Wong, Ester Calvo Fernández, Evan Paull, Sunny Jones, Junqiang Wang, Heeju Noh, Beatrice Salvatori, Alejandro Chavez, Andrea Califano

## Abstract

Pooled CRISPRi-mediated silencing of >1,000 transcriptional regulators expressed in single colorectal adenocarcinoma cells, followed by single-cell RNA-seq profiling at two timepoints, 1 day and 4 days, allowed reverse engineering the underlying tumor context-specific, causal regulatory network. Furthermore, the availability of experimentally derived, highly multiplexed gene reporter assays for each regulator, as identified by this analysis, allowed accurate assessment of differential protein activity following silencing of each regulator, thus providing proof-of-concept for generating comprehensive, tissue-specific networks of transcriptional and post-translational interactions. Analysis of this causal network allowed elucidation of complex autoregulatory mechanisms that have eluded previous computational approaches and supported systematic elucidation of cooperative mechanisms, where one regulatory protein can modulate the activity of another regulatory protein, as well as transcriptional mimicry, where one regulatory protein can phenocopy others.

## Introduction

Precise control of transcriptional cell state by transcription factors and co-factors is critically required across all aspects of mammalian cell pathophysiology: from determining cell identity during lineage differentiation to providing mechanistic control of cancer cell plasticity. Yet, while global regulatory mechanisms involved in lineage specification have been experimentally elucidated in a few model organisms, such as *C. elegans* (Murray 2018) and sea urchin (Davidson, Rast et al. 2002), those relevant to pathophysiology of mammalian cells are only sparsely understood, mostly as a result of small-scale, hypothesis-driven assays (Takahashi 2017), or from sparsely validated computational reverse engineering algorithms (Basso, Margolin et al. 2005, Huynh-Thu, Irrthum et al. 2010). A key limitation of existing gene regulatory network (GRN) models is their inability to resolve the directionality (i.e., *causality*) of transcriptional interactions, thus limiting our ability to elucidate complex autoregulatory logic controlling cellular homeostasis, in both physiologic and pathologic cell states. Homeostatic control of cell state represents one of the most critical functions of GRNs, which is directly responsible for the adaptive behavior necessary to maintaining cellular identity and function in normal cell physiology as well as for the cell adaptation mechanisms that allow disease-related cells to adapt and escape pharmacological treatment. Indeed, elucidation of autoregulatory loops has been the Achilles’ heel of virtually all proposed computational approaches and has required complex, multi-gene experimental assays for identification and validation, even on a relatively small scale (Rajbhandari, Lopez et al. 2018).

Moreover, we and others have shown that the availability of accurate and comprehensive GRN models is critical for the utilization of network-based algorithms for the study of cellular phenotypes and functions. These include methodologies for (a) accurately measuring protein activity from RNA-seq profiles (Alvarez, Shen et al. 2016), including at the single-cell level (Obradovic, Chowdhury et al. 2021), (b) elucidating key regulators of cellular programs (Boorsma, Lu et al. 2008, Alvarez, Shen et al. 2016, Aibar, Gonzalez-Blas et al. 2017), (c) elucidating Master Regulators representing mechanistic determinants of cell state in cancer and other diseases (Carro, Lim et al. 2010, Aytes, Mitrofanova et al. 2014, Rajbhandari, Lopez et al. 2018), (d) predicting proteins capable of reprogramming cell-state (Dutta, Le Magnen et al. 2016, Arumugam, Shin et al. 2020), (e) dissecting drug Mechanism of Action (MoA) (Woo, Shimoni et al. 2015).

Unfortunately, since most GRNs have been computationally dissected (Basso, Margolin et al. 2005, Pe’er 2005, Wang, Saito et al. 2009) and only sparsely validated, there are still lingering concerns about whether they may truly recapitulate the underlying regulatory logic of the cell. Finally, virtually all computationally dissected GRNs have been reconstructed from steady-state molecular profiles - i.e., representative of cells whose dynamics are slow compared to the half-life of gene products. These GRNs will thus, by definition, miss key elements of dynamic/time-dependent gene regulation, including the ability to model autoregulatory loop-mediated oscillatory behavior following exogenous and endogenous perturbations (Ma, Wagner et al. 2005).

Unfortunately, in contrast to the experimental elucidation of protein-protein interaction networks (Rual, Venkatesan et al. 2005), the development of genome-wide experimental technologies for the reverse engineering of GRNs is challenging and remains elusive. Ideally, one would perturb each regulatory protein and assess its effect on the expression and activity of all other genes and proteins they encode, respectively, at multiple timepoints. While this would have been essentially unfeasible, due to cost and effort, until now, we were able to combine several recently-developed technologies into a fully integrated experimental pipeline for the systematic, single-cell-based dissection of GRNs. This Transcriptional Regulator Knockdown (TReK) pipeline leverages highly-multiplexed, pooled CRISPRi-mediated silencing of transcriptional regulators in single cells, followed by single-cell RNA sequencing (scRNA-seq) at multiple timepoints, including a short timepoint (24h) to capture transient regulatory interactions and a longer timepoint (96h) to capture regulatory interactions as cells start to achieve steady-states. While still coarse, we expect this to provide an initial framework to start revealing time-dependent regulatory cascades controlling critical genetic programs (Figure 1A).

**Figure 1.** Conceptual overview of study and summary of dataset. (A) Concept of capturing dynamic transcriptional targets by early timepoint data collection. (B) Computational flowchart. Raw input data (red) was processed to directly generate biological readout (blue) or with the help of transcriptional network/regulons (yellow) construction. (C) Timeline for sgRNA transduction, selection, and doxycycline-induction. Lentiviral sgRNA library including >3,000 sgRNA species was transduced into colorectal cancer cell line HT-29 pre-transduced with constitutive dCas9 vector. Puromycin (8 ug/ml x 3 days) was added 24 hours post-transduction to select for sgRNA-positive cells. T2 sample was directly collected after 3 days of puromycin selection. For T1 experiment, HT-29 was pre-transduced and selected for the inducible dCas9 vector. At the time of the experiment, 1ug/ml doxycycline was added and cells were collected after 1 day of doxycycline induction. (D) Important metrics across two timepoints. At both timepoints, we profiled CRISPRi-perturbed transcriptomes in 70k to 80k single cells. Around 95% of these cells generated a high amount of UMI counts and 60-70% of the cells had one unique sgRNA detected. Knockdown efficacy for each TR was estimated based on the most efficient sgRNA among all 3 sgRNAs targeting that particular TR. (E) Cell coverage per sgRNA perturbation in T2 experiment. Only single cells with a unique sgRNA and more than 5,000 total UMI counts are considered.

To achieve optimal CRISPRi-mediated silencing of target genes, we combined an enhanced CRISPRi repressor (Yeo, Chavez et al. 2018) with improved single guide RNA (sgRNA) predictions (Sanson, Hanna et al. 2018) into pooled, single-cell CRISPRi screens with transcriptomic readout using the recently developed CROP-seq technology (Datlinger, Rendeiro et al. 2017). In addition, we developed an inducible version of Yeo et al.’s enhanced CRISPR repressor (Yeo, Chavez et al. 2018), thus allowing precise timing of cell harvesting and scRNA-seq profiling as early as 12h after repressor induction and gene silencing.

As a proof of concept, we applied TReK pipeline to dissect colorectal cancer-specific GRNs, via single-cell perturbation and scRNA-seq profiling of HT-29 cells. Specifically, we generated single-cell profiles of >150,000 HT-29 cells at two timepoints (24h and 96h), following pooled, CRISPRi-mediated, single-cell silencing of 1,007 transcriptional regulators, thus measuring 6 billion unique molecules (UMI-gene pairs), while dramatically reducing labor, assay costs, and potential batch effects. The genes silenced in this study were selected to include virtually all actively transcribed transcriptional regulators (hereafter TRs) - including transcription factors (TFs) and co-factors (co-TFs), such as (de)acetylases, (de)methylases, and other chromatin remodeling enzymes - that were detected as either significantly expressed or transcriptionally active in this cell line. Colorectal adenocarcinoma (COAD) was selected as a relevant tumor context because of lack of effective therapy and recent elucidation of Master Regulator proteins defining 8 molecularly distinct subtypes, including 3 subtypes highly enriched in MSI-high tumors and an aggressive, genomically-stable subtype characterized by very poor outcome (Paull, Aytes et al. 2021). Thus, we expected that a high-quality experimentally dissected GRN would be a first step towards the elucidation of critical COAD tumorigenesis and progression mechanisms.

When integrated into existing computational reverse engineering methods, the TReK data was instrumental in allowing significant improvement in GRN model accuracy, as objected assessed based on the ability to recapitulate changes in the activity of the proteins encoded by silenced genes both at 24h and 96h. These studies allowed reconstructing many established transcriptional and post-translational mechanisms, refining our understanding of protein-protein interactions in complexes, and most critically, reconstructing fully integrated, causal GRNs that include both transcriptional and post-translational interactions (Figure 1B).

## Results

### Large-scale CRISPRi-mediated TR silencing in HT-29 cells at two timepoints

To generate the perturbational data needed for the systematic dissection of COAD-specific molecular interactions, leading to a full GRN, we used assays that combine CRISPRi (Gilbert, Horlbeck et al. 2014) and CROP-seq (Datlinger, Rendeiro et al. 2017) technologies (Figure S1; Methods). Specifically, we measured changes in gene expression, within individual HT-29 cells, as measured by Chromium scRNA-seq profiles, following pooled CRISPR/dCas9-mediated silencing of each significantly expressed or active transcriptional regulator protein (see Methods) (Figure 1C).

Using this pipeline, we thus generated scRNA-seq profiles following silencing of 1,007 genes in HT-29 cells, at two timepoints (T1 = 24h, T2 = 96h), using three sgRNAs per gene. To select target genes, we first leveraged a manually curated repertoire of 2,527 transcriptional regulators (hereafter TRs) annotated as either TFs or co-TFs in Gene Ontology (Alvarez, Shen et al. 2016) and then sub-selected those that were either significantly expressed or active in HT-29 cells (see Methods). Thirty scrambled sgRNAs (sgCtrls) were also included as non-targeting negative controls, resulting in a library comprising 3,051 sgRNAs (Table S2). To achieve robust CRISPRi-mediated gene silencing, we engineered lentiviral plasmids supporting either constitutive or doxycycline-inducible expression of the enhanced repressor dCas9-KRAB-MeCP2 (Yeo, Chavez et al. 2018) (Figures S2A and S2B; Methods). To assess the actual length of effective gene silencing, we assessed the time required for the CRISPRi repressor to be transcribed and translated, following its doxycycline-inducible expression (Figure S2B; Methods). This shows that cells collected at 24h following dCas9-KRAB-MeCP2 induction (T1), have effectively undergone ∼12h of sustained repression. For the second timepoint, we used cells transduced with a constitutive dCas9-KRAB-MeCP2 expression plasmid (Figure S2C), harvested at 96h following lentivirally-mediated transduction with the sgRNA library (T2). HT-29 cells were selected also because, in mid-log growth, they have high transcriptome complexity, with a median of 35,000-42,000 UMIs/cell, at about 30% sequencing saturation and >90% of reads in cells, thus supporting follow-up analyses.

At each timepoint, >70,000 single-cell transcriptomes were sequenced (Figure 1D), followed by removal of low-quality profiles (<5,000 UMIs) and of profiles with two or more detected sgRNAs. More than 60% of the single cells passed this strict QC filter, thus producing a dataset with a median of 14-15 high-quality single-cell transcriptomes for each sgRNA (Figure 1E), yielding a total of 44-47 cells, on average, in which each target gene had been silenced by one of the three corresponding sgRNAs.

Achieving efficient, systematic gene silencing across a large pooled gene library is still a rather elusive goal with first-generation CRISPRi. Thus, it was critical to validate the performance of our second-generation CRISPRi implementation (Sanson, Hanna et al. 2018, Yeo, Chavez et al. 2018). To accomplish this, we performed a careful quality-control analysis. We analyzed the efficacy of gene silencing compared to controls (Figure S4), measured the correlation between different gRNAs targeting the same gene (Figures S5A and S5B), and compared our scRNA-seq gene silencing to published CRISPR knockout data (Figures S5G and S5H). In addition, we analyzed the factors that affect successful silencing (Figures S5C-S5F; Methods). Overall, we found that for 70% of tested TRs, we could achieve 50% silencing, sufficient to reliably identify targets, at both timepoints, with the later timepoint providing critical information for proteins with a long half-life.

### Reconstruction of GRNs inferred from computational, experimental, and integrative analysis

As previously discussed, computationally inferred GRNs are typically reverse engineered from steady-state data, thus potentially reflecting broad co-expression patterns that may not be representative of direct regulatory events. Moreover-with the exception of Dynamic Bayesian Networks (Smith, Yu et al. 2006), which have never been validated in a mammalian context-, these methodologies are unable to elucidate auto-regulatory control mechanisms and fail to adequately represent regulators proteins whose expression represents an inadequate proxy of transcriptional activity.

For instance, while the ARACNe algorithm (Margolin, Nemenman et al. 2006) effectively eliminates indirect interactions using the Data Processing Inequality theorem and has been extensively experimentally validated, it still suffers from the above-mentioned limitations. To objectively assess the ability of TReK assays to improve GRN reconstruction quality, we compared three networks, including (a) purely computationally-inferred ARACNe networks inferred from the TReK scRNA-seq profiles at T1 or T2, (b) a purely experimental network assembled from genes that are differentially expressed following silencing of each regulator’s silencing at T1 or T2, and (c) an integrated network that combines both computational and experimental analyses (see Methods). For the latter, we modified a two-branch generalized linear model (GLM) that was previously used for single-cell differential expression analysis (Finak, McDavid et al. 2015) (Figure 2A; Methods). The integrated network combines ARACNe p-values for individual regulator-target interactions as well as the differential expression of the target following silencing of the regulator in the TReK assays, thus accounting for both interaction directionality and strength. Following GLM analysis, 65% and 61% of the TR targets inferred at T1 and T2 were derived from TReK data, respectively, with the remaining 35% and 39% coming from ARACNe.

**Figure 2.** Gene regulatory network reconstructed by CRISPRi perturbation data. (A) Regulatory coefficient matrices on continuous space or binary space were fitted to the single-cell normalized expression matrix or binarized expression matrix given the target TR perturbation and other covariates in the design matrix. Coefficient matrices were subsequently consolidated and converted to VIPER-compatible regulons. (B) Barplot showing fraction of significantly inactivated (*p* < 0.05) TRs inferred by different networks. For each TR, protein activities were inferred in each perturbed single cell and then integrated by Stouffer’s method. (C) Correlation between perturbed protein activities inferred by integrated network and number of target genes. Each dot represents a TR. The number of target genes is represented by the number of differentially expressed genes (DEGs) after the TR perturbation. (D) Correlation between perturbed protein activities and “importance scores”. Each dot represents a TR. “Importance score” of each TR is represented by its protein activity in HT-29 comparing to other CCLE cell lines, as calculated in Methods. The integrated network was used in HT-29 VIPER analysis. (E) HT-29-specific DNase-Seq peaks are obtained from Cistrome database (Liu, Ortiz et al. 2011) as described in Methods. Statistical significance was calculated using the one-tailed Mann-Whitney U test.

### Integrative GRN reverse engineering outperforms both computational and experimental methodologies

To assess network quality, we combined two metrics including (a) detection of regulatory protein activity decrease in the subset of cells where they were effectively silenced, (b) consistency with DNase-seq data (Mei, Qin et al. 2017, Zheng, Wan et al. 2019). For the first metric, we assessed protein activity using the VIPER algorithm, as determined by the differential expression of its target genes as represented in each of the three networks (computational, experimental, and integrated). A highly accurate network would show gene expression increase of induced targets and decrease of repressed targets in cells where the regulator is silenced. Thus, the more accurate the network is, the more accurately the activity of a TR should be measured by the differential expression of its regulon genes.

For this analysis, we selected 250 TRs, whose CRISPRi-mediated silencing produced ≥ 50% silencing at both timepoints, in at least 15 cells/sgRNA (see Methods), as the gold standard set, based on the assumption that their transcriptional activity would be significantly affected by silencing and presented in a sufficient number of cells to support the analysis. We then assessed GRN accuracy by 5-fold cross-validation within each timepoint (see Methods), based on VIPER-based TR inactivation assessment based on the differential gene expression signature of cells where the TR was effectively silenced vs. negative control cells transduced with non-targeting sgRNAs. In addition, we also performed cross-timepoint validation—i.e., assessing how well GRNs inferred at one timepoint could improve protein activity inference accuracy at the other timepoint (see Methods). The analysis clearly shows that while the experimental GRN outperforms the computational one, especially at the shorter timepoint (24h) the integrated GRN significantly outperforms both (Figure 2B). These results are consistent with expectations that computational GRNs will perform better as cells achieve steady-state and will thus fail to identify transient regulatory events that may be compensated by autoregulatory loops. It also suggests that the pure experimental data may be noisy and thus introduce false positive and false negative targets that are then optimally refined by the computational analysis.

To understand the factors that most contributed to improved GRN performance, we first tested whether TRs inducing a larger signature of significantly differentially expressed genes would outperform those with smaller effect size. Indeed, inferred protein activity accuracy was significantly correlated with the number of differentially expressed genes (*p* < 2.7e-9) (Figure 2C). We also tested whether silencing the most likely Master Regulators of cell-state would induce stronger transcriptional signatures. Specifically, we first inferred the differential activity of all tested TRs in HT-29 against the average of all CCLE cell lines (Ghandi, Huang et al. 2019) using VIPER with the integrated network (see Methods). We then measured the correlation between their VIPER-inferred differential activity in HT-29 cells and their differential activity following CRISPRi-mediated silencing. Correlations were significant (*p* = 2.4e-3 at T1; *p* = 4.1e-9 at T2) suggesting that silencing critical regulators of cell-state would induce stronger transcriptional signatures (Figure 2D). Finally, across 4 cross-validations, 27 of the 38 TRs (71%) that were silenced in ≥60 cells were significantly inactivated based on VIPER analysis (Figures 2C and 2D), suggesting that TR targets inference—and thus activity measurement—could be further improved by increasing the number of cell profiles in TReK assays. Indeed, saturation analysis shows that the VIPER-measured protein activity of most TRs sampled in ≥ 60 cells converged to a narrow range as an increasing number of cells were included in the analysis (Figure S6).

For the second metric, we show that the consistency of target gene genomic loci with open chromatin, as assessed by DNase-seq in HT-29 cell lines (Mei, Qin et al. 2017, Zheng, Wan et al. 2019), significantly (*p* = 1.52e-8 at T1; *p* = 3.43e-7 at T2) improves in the integrated GRN at both timepoints, compared to the ARACNe computational networks (Figure 2E).

Based on these results, the integrated computational/experimental GRNs was selected as the optimal one in all subsequent analyses.

### Integrated GRNs capture multi-layer molecular interactions

A key goal of cellular network reverse engineering has been the assembly of molecular interaction networks that integrate distinct mechanisms of gene-product interaction. GRNs capture transcriptional regulation effects, mediated by regulatory proteins binding the regulatory region of a target gene, either directly or via cognate binding partners. However, these networks fail to capture the post-transcriptional effect of protein X on protein Y. For instance, two transcription factors may bind the same DNA-binding motif (e.g., ARNT and MYC binding E-boxes) thus inducing a stoichiometric mediated dependency where over/under-expression of one (e.g., ARNT) may induce activity increase/reduction of the latter (i.e., MYC), respectively (Wang, Saito et al. 2009). Similarly, a chromatin remodeling enzyme may result in epigenetic silencing of the targets of another TR. Finally, a TR may regulate a ubiquitin ligase adapter protein, thus inducing proteasomally-mediated degradation of another TR (Chen, Alvarez et al. 2014). A unique advantage of the TReK data is that it supports dissection of both transcriptional and post-transcriptional interactions, based on the VIPER inferred activity of each TR, following silencing of another TR, allowing reverse engineering of the first integrated, multi-layer TRN (transcriptional regulatory network) and ARN (activity regulatory network) for a human cell.

To further improve sensitivity, we integrated the T1 and T2 ARN networks, by taking the union of the regulons inferred at each timepoint, while removing mode-inverted interactions at T2—i.e., interactions detected as positive at T1 and repressive at T2, or vice versa (see Methods). This produced an integrated ARN comprising 23,607 activity-regulating interactions (false discovery rate, FDR < 0.1) involving 949 of the 1007 TRs. Since protein activity can be regulated by many complementary mechanisms, we studied several TR interaction mechanisms in greater detail. Specifically, we considered four non-mutually exclusive mechanisms that may account for ARN interactions, including transcriptional regulation, direct or indirect relationships in the same pathway, physical protein-protein interactions, and co-regulation of the same target genes.

To assess transcriptionally mediated ARN interactions—i.e., interactions where silencing of TR*x* affects the activity of TR*y* because TR*x* transcriptionally regulates TR*y*, we assessed how much of the differential activity of each TR could be accounted for by the differential expression of its encoding gene. Overall, only 2.9% (*n* = 685) of inferred ARN interactions could be accounted for by this mechanism (Figures 3A and 3B). Second, an examination of each TR’s pathway membership showed that ∼44% (*n* = 10,347) of ARN interactions were detected between same-pathway TRs. Third, ∼7.0% (*n* = 1,650) of ARN interactions were between structural cognate binding partners, as reported in protein-protein interaction databases (see Methods). Finally, a small fraction (4.2%, *n* = 980) of the interactions were explained by significant overlap between regulons (see Methods). In addition to ARN interactions, the TRN comprised *n* = 121,414 interactions (FDR < 0.2) detected by differential expression analysis, including *n* = 9,812 TR-TR interactions.

**Figure 3.** TReK multi-layer regulatory network. (A) Mechanisms contributing to edges in the activity regulatory network. Multiple analyses and datasets were used to characterize each edge (Methods). The fraction of edges annotated by each mechanism is shown in bars. The cumulative fraction of all edges with at least one mechanism is shown by the line. Transcriptional, characterized as transcriptional by the RPT analysis. PPI, protein-protein interaction. Regulon, overlapping regulon. (B) Overlap of edge mechanistic characterizations that are summarized in (A). (C,D) Heatmaps representing gene expression (C) and protein activity (D) profiles after perturbation of Wnt-TGF pathway regulators. Statistical insignificant values (*p* > 0.05) are set to be white. (E) Multi-layer regulatory network represented by (C,D). If both transcriptional and activity-level regulation are present, only transcriptional regulation is shown. Essential genes are defined as having Achilles dependency scores higher than 0.9. (F,G) Distribution of different patterns of 2-gene feedback loops in multi-level regulatory networks. (H,I) Distribution of different patterns of 3-gene feedback loops in multi-level regulatory networks. (J) Number of 2-gene or 3-gene feedback loops participated by different TRs. TRs were grouped into quintiles by their VIPER-inferred protein activities in HT-29 against all other CCLE cell lines. Integrated networks were used for inferring protein activities.

### Analysis of integrated TRN/ARN networks

Integration of TRN and ARN interactions into a multi-layer network (MLN = TRN + ARN) contributes to a more holistic understanding of biological mechanisms. For instance, we used the more comprehensive T2 network to analyze an MLN sub-network comprising *MYC* and six established MYC regulators in the Wnt and TGF-β pathways (Figures 3C-3E). Both pathways are known to play critical roles in COAD, contributing to increased activity of the MYC proto-oncogene, by repression of TGF-β signaling and hyperactivation of Wnt signaling (Cancer Genome Atlas 2012). Our analysis recapitulated key, well-established interactions, such as the reciprocal activation (positive feedback loop) between *TCF7L2* and *CTNNB1* (Brantjes, Barker et al. 2002), which is a crucial step in the activation of the Wnt signaling pathway. Notably, the essential proto-oncogenes (*MYC*, *CTNNB1*, and *TCF7L2*) form a highly interconnected module with multiple activating interactions (Figure 3E, red edges). In contrast, the non-essential tumor suppressors *SMAD2* and *SMAD3* are identified as repressors of the MYC-centric oncogenic module (Figure 3E, blue edges). While some of these interactions are well-established in the literature (Yagi, Furuhashi et al. 2002, Rennoll and Yochum 2015), our analysis identified several novel interactions, such as the *MYC*-mediated transcriptional activation of *TCF7L2* and *CTNNB1*, and the negative feedback loops formed by *SOX9* with *MYC*, *CTNNB1*, and *TCF7L2*, which further increase our understanding of TGF-β/Wnt-mediated MYC dysregulation in COAD. Not surprisingly, consistent with the fact that *SMAD4* has a nonsense mutation in HT-29, according to the COSMIC database (Tate, Bamford et al. 2019), we did not detect SMAD4 interactions with other proteins.

Analysis of the full MLN identified a preponderance of ARN vs. TRN TR interactions (*n* = 23,607 vs. *n* = 9,812, respectively) (Figure S7A). To further delineate transcriptional and post-transcriptional regulation in the ARN, we removed ARN interactions that were explained by a corresponding TRN interaction, this process resulted in *n* = 22,922 post-transcriptional interactions, including activating vs. repressive TR interactions (*n* = 20,278 vs. 12,456, respectively) (Figure S7B). The resulting networks showed statistically significant overlap with both MultiNet (*p* = 2.94e-19) (Khurana, Fu et al. 2013) and HumanNet (*p* = 3.86e-60) (Hwang, Kim et al. 2019) (Table S3).

### Auto-regulatory loops are significantly over-represented in the multi-layer network

Auto-regulatory feedback represents a ubiquitous regulatory mechanism in biological systems, and some associated feedback loop motifs are significantly overrepresented (Alon 2007). A unique aspect of the TReK assays is that they allow direct, causal dissection of autoregulatory loops because each TR is independently silenced, thus breaking the upstream but not the downstream logic of the loops in which it is involved.

#### 2-Protein Loops

Systematic analysis of feedback loops comprising 2 proteins in the MLN identified 1,816 instances, comprising 3 different motifs: positive feedback (two positive interactions), negative feedback (one positive and one repressive interaction), and genetic switches (two repressive interactions) (Figure 3F). Compared to the null distribution obtained by randomizing MLN edges, while preserving its overall topology, we observed significant overrepresentation of all loop motifs (3.3x, 2.4x, 1.89x) (Figure 3G). Interestingly, compared to the total number of 2-protein loops, positive feedback loops were further over-represented, while negative feedback loops and genetic switches were slightly under-represented (*p* < 2.2e-16 by chi-square test).

#### 3-Protein Loops

In addition, 10,260 3-protein loops were identified across 4 different motifs (Figure 3H), depending on the number of positive and negative interactions (Figure 3I). Similar to 2-protein loops, we observed overrepresentation of positive loops (*p* < 8.94e-12 by chi-square test).

Loops comprising proteins with greater differential activity in HT-29 cells were more likely to be detected. Indeed, our analysis showed that loops comprising proteins in the first quintile of all TRs, ranked based on their differential activity in HT-29 against the average of CCLE, formed almost twice as many loops compared to other proteins (Figure 3J). This suggests that the proteins responsible for implementing and maintaining cell-state homeostasis are more likely to participate in feedback loops.

### Variable knockdown efficacy and time-course data reveal systematic use of autoregulatory feedback

While variable sgRNA silencing efficacy is normally seen as a potential limitation of the CRISPRi technology, in the context of GRN reverse engineering, it can actually help further elucidate the presence of regulatory feedback (auto-regulation). In the absence of auto-regulatory feedback, the more effective the silencing of a TR, the stronger the activation or repression of its targets. However, auto-regulatory loops may induce significant, time-dependent oscillations in regulatory targets expression or activity.

For instance, of the 3 sgRNAs targeting *MYC*, two induced moderate (60%) and weak (17%) silencing, respectively, while the third failed to produce any silencing. Gene Set Enrichment Analysis (GSEA) (Subramanian, Tamayo et al. 2005) confirmed that enrichment of MYC targets in genes under-expressed following its silencing tracked with silencing efficacy (Figures 4A-4C). Similar correlation between silencing efficacy and target gene expression in the associated pathways was observed for many other genes, such as *CTNNB1*/Wnt and *MYBL2*/cell cycle (Figure S8). More interestingly, however, for some TRs, we observed virtual inversion of gene expression signature behavior as a function of sgRNA efficacy, implicating the existence of strong autoregulatory mechanisms. E2F7, a transcriptional repressor of the E2F family (de Bruin, Maiti et al. 2003, Di Stefano, Jensen et al. 2003, Westendorp, Mokry et al. 2012), provides an interesting example of such behavior. Under normal circumstances, repressing E2F7 activity will cause upregulation of its target genes (Segeren, van Rijnberk et al. 2020). Indeed, three sgRNAs targeting *E2F7* induced significant E2F target upregulation (*p* = 0.01, 0.037, and 0.23, respectively) at the earlier timepoint (T1) (Figures S9A-S9C). However, at the later timepoint (T2 = 96h) the guide with the highest efficacy induced complete inversion of target expression (mode inversion), with significant E2F target downregulation (*p* < 0.001) (Figures 4D, 4E, and S9D). Since E2F targets, which are involved in cell cycle progression, are known to undergo oscillatory behavior (Westendorp, Mokry et al. 2012), consistent with overcompensation and mode inversion following its most significant silencing. Similar to E2F7, we observed highest-efficacy vs. low-efficacy mode inversion for 57 additional TRs at T2, suggesting their participation in strong autoregulatory loops. For 10 (>17.5%) of those highest-efficacy sgRNAs we observed mode inversion between T1 and T2 (Table S4). This is 7-fold what would be expected in the null hypothesis (see Methods).

**Figure 4.** Different modes of regulation revealed by different patterns of transcriptome responses. (A-C) Correlation between *MYC* perturbation efficacy and differential expression enrichment of MYC targets. (A) Log(CPM+1) of *MYC* expressed in cells expressing sgRNA targeting a different gene (“All Other”), cells expressing scrambled sgRNAs (“NonTargeting”), and cells expressing *MYC*-targeting sgRNAs shown in violin-boxplots. (B) Gene set enrichment analysis of MYC targets on gene expression signatures after 3 different *MYC*-targeting sgRNA perturbations. Empirical p-values are shown. (C) Correlation between *MYC* perturbation efficacy and differential expression enrichment of MYC targets. Since knockdown efficacy was calculated against 100 randomly sampled pseudo-bulk control samples (see Methods), the median ± SD for all 100 knockdown efficacy values are shown as horizontal error bars. Each GSEA analysis was run 5 times and the median ± SD of 5 normalized enrichment scores (NES) are shown as vertical error bars. (D) Log(CPM+1) of *E2F7* expressed in different groups of cells. (E) Correlation between *E2F7* perturbation efficacy and enrichment of E2F targets at both timepoints. (F) Mode of regulation of Wnt-TGF pathway regulators on hallmark cellular processes. Specific regulations that are corroborated by published evidence are listed in Table S5. Summary across each row or column is shown as stacked histograms. Results from T2 experiment are shown.

### Regulation of Hallmark Cellular Processes

To elucidate regulators of fundamental cellular processes, as cell achieve stable state, we assessed enrichment of genes differentially expressed at 96h following silencing of each TR in genes comprising 50 “hallmark” cellular processes by GSEA analysis (Liberzon, Subramanian et al. 2011, Liberzon, Birger et al. 2015). Potential mode of regulation was categorized as activation if silencing a TR caused downregulation of a cellular process, repression if silencing a TR caused upregulation of a cellular process. To address the effect of potential auto-regulatory loops (see previous section), we included a third category called “mixed” if both downregulation and upregulation were observed after silencing a TR with different sgRNAs (see Methods). To minimize false discovery rates, we only considered TRs for which at least 2 sgRNAs produced concordant enrichment. While this undoubtedly prevents detection of bona fide hallmark process regulators, for which there was only 1 effective sgRNA, the analysis revealed many COAD-specific hallmark process regulators, including both established and previously uncharacterized ones. For example, considering 16 TRs in the Wnt/TGF pathway (Figure 4F), we recapitulated regulation of both mTORC1 signaling and DNA repair by MYC (Hironaka, Factor et al. 2003, Karlsson, Deb-Basu et al. 2003, Yue, Jiang et al. 2017), of spermatogenesis by ID2 (Sablitzky, Moore et al. 1998), and of epithelial-mesenchymal-transition (EMT) by β-catenin (Vu and Datta 2017). Indeed, cursory search of the literature supported 24 of the 55 inferred hallmark regulatory interactions associated with the 16 TRs (Table S5). Thus, availability of this analysis to all 1,007 TRs provides a powerful new resource to study regulatory processes and mechanisms in this COAD (Table S6).

### Identification of upstream regulators and co-factors

The TReK assays also support identification of regulatory hierarchies and co-transcriptional factors. For instance, as expected, CRISPRi-mediated silencing of MAX, an established co-factor and modulator of Myc activity (Dang 2012), significantly affected MYC target expression (*p* < 0.001) (Figures S10A-S10D). To further confirm enrichment of potential MYC modulators in TRs that affect expression of MYC tatgets, we first assessed all TRs whose silencing affected the targets of a second TR, using the CINDy algorithm (Giorgi, Lopez et al. 2014), CINDy is an improvement of the original MINDy algorithm (Wang, Saito et al. 2009), which uses the conditional mutual information *I[TR;T|M]* between a *TR* and its targets *T*, given the expression of a candidate modulator *M* to predict TR activity modulators. Experimental validation rates for these algorithms have been in the 70%- 80% range and they have been shown to recapitulate 60% to 70% of known upstream pathway interactions in lung adenocarcinoma and lymphoma (Wang, Saito et al. 2009, Giorgi, Lopez et al. 2014).

We found significant overlap between CINDY and TReK-based predictions of candidate MYC-activity modulators (Figure S10E), with 115 of the 289 TReK-predicted ones also predicted by CINDY (*p* = 0.014 by Fisher’s exact test). MLN-based characterization of these 115 TRs in the MLN identified 5 as upstream MYC transcriptional regulators, 13 as upstream post-transcriptional regulators, and 27 as physical interactors, based on PrePPI protein-protein interactions (Zhang, Petrey et al. 2013) (Figure S10F).

## Discussion

While experimental and computational methodologies for the dissection of protein-protein interaction (PPI) networks have achieved relative maturity (Zhang, Petrey et al. 2012, Rolland, Tasan et al. 2014), the inference of transcriptional and post-transcriptional interactions is still broadly debated, despite the role they have played in elucidating novel biological mechanisms, disease drivers, and phenotype (Chen, Alvarez et al. 2014, Alvarez, Shen et al. 2016, Rajbhandari, Lopez et al. 2018, Arumugam, Shin et al. 2020). Among others, current limitations include: (a) the relatively low value of DNA-binding motifs, given their ubiquitous presence in open chromatin regions, the fact that most TFs do not bind DNA via their canonical motif, and the uncertainty of whether binding induces functional regulation (b) the inability of computational methods to assess causality and directionality thus preventing detection of autoregulatory loops playing a key role in cell homeostasis and (c) the very sparse experimental validation of transcriptional targets. This is largely due to the fact that technologies for the systematic RNA-seq profiling of specific cellular contexts following perturbation of each transcriptional regulator (TR) have been lacking or have been excessively labor-intensive and costly.

By combining two technologies, including 2^nd^ generation, inducible, CRISPRi-mediated gene silencing and CROP-Seq, supporting single-cell RNA-seq profiling with knowledge of the specific single guide RNA(s) in each cell, we have developed a Transcriptional Regulator Knockdown methodology (TReK) for the systematic, experimentally-based reconstruction of gene regulatory (GRN) and activity regulatory (ARN) networks. GRNs comprise molecular interactions where a specific TR affects the expression of another gene, while ARNs comprise molecular interactions where a specific TR affects the transcriptional activity of another TR.

While the current implementation of TReK provides a critical proof-of-concept of the technology, its application to a colorectal cell line HT-29 already overcomes critical limitations of current methodologies for GRN inference, while allowing construction of previously unavailable ARNs representing key, biologically relevant post-transcriptional and post-translational interactions. In particular, TReK provides key experimental evidence supporting TR target inference based on changes in their expression following CRISPRi-mediated TR silencing at two timepoints. The rationale for the two timepoints is multifold. First, this allows incorporating proteins with different half-lives in the analysis. Second, this allows detection of autoregulatory effects where the targets of a TR are inversely correlated in their expression at the two timepoints; critically, decoupling TRs from their upstream regulation, via direct biochemical perturbation, allows deconvoluting both interaction directionality (i.e., A→B vs. A←B) and autoregulatory loops (e.g., A→B→A), which has been challenging using computational approaches. The data presented here, which include analysis of >150,000 single-cell profiles representing CRISPRi-mediated silencing of 1,007 expressed TRs in HT-29 cells at 24h and 96h, not only confirms previous interactions, such as those between MYC and WNT/β-catenin pathway proteins in colon cancer, but identifies thousands of novel interactions and experimentally assessed effects of individual TRs on cancer hallmarks, including >15,000 experimentally dissected 2- and 3-protein loops contributing to cellular homeostasis.

Critically, while still limited, this study provides key indications on the number of cells that will be necessary for future implementations of the TReK assays to ensure accurate representation of all transcriptional regulators. Specifically, saturation analysis (Figure S6) suggests that ≥ 60 cells per silenced TR would be sufficient to stabilize protein activity predictions based on TR targets, using the VIPER algorithm, thus opening the way to dramatic improvements in the assessment of mechanistic determinants of cellular phenotypes, not only in cancer but across all accessible cell states. In addition, similar studies targeting signaling proteins would also be possible, allowing the activity of TRs to be used as gene reporter assays for their upstream signaling modulators as well as for deconvoluting the complete logic of signal transduction, including autoregulatory loops. The cost of a TReK experiment is still relatively high; however, the results of this proof-of-concept experiment combined with the advent of additional technologies such as scifi-RNA-seq (Datlinger, Rendeiro et al. 2021) and the ever-decreasing cost of sequencing suggests that a full TReK experiment including all TRs and signaling proteins (∼6,500 proteins in total) at 4 timepoints and with an average of 60 cells per silenced protein, could be performed for less than $20,000, thus providing an entirely novel perspective on transcriptional and post-translational regulation. Furthermore, inclusion of proteins representing targets of FDA-approved or late-stage development small molecular compounds, combined with their genome-wide perturbational profiles (Woo, Shimoni et al. 2015), could provide a remarkable ability to deconvolute drug mechanism of action and to identify drugs capable of activating or inactivating any desired regulatory program, as previously shown using computationally based GRNs, only with dramatically improved accuracy and sensitivity.

Among the limitations of the current approach, one should consider that perturbations may be implemented in single cells in different stages of cell cycle. As a result, cell cycle-dependent regulatory events may require additional regression of cell cycle stage. However, with ≥ 60 cells per silenced TR, such regression would become eminently possible. Additionally, current sgRNAs present highly different gene silencing efficiency, with about 10% to 20% of the genes not silenced at all. However, this can be effectively addressed by performing a smaller-scale TReK experiment to first select sgRNAs that provide optimal silencing, thus saving significant effort and cost in the larger experiments by removing the need to sequence a large number of cells where the target gene is not effectively silenced.

Analysis of the GRN and ARN networks already produced key novel findings. However, much as it has happened for PPI networks, comprehensive assessment of the TReK GRN/ARN networks will only be possible as they are explored by biologists with precise, phenotypically driven questions.

## Acknowledgements

This work was supported by an NCI Outstanding Investigator Award (R35 CA197745) and two NIH Shared Instrumentation Grants (S10 OD012351 and S1 0OD021764), all to AC. Also, this research was funded in part through the NCI Cancer Center Support Grant (P30 CA013696).

## Methods

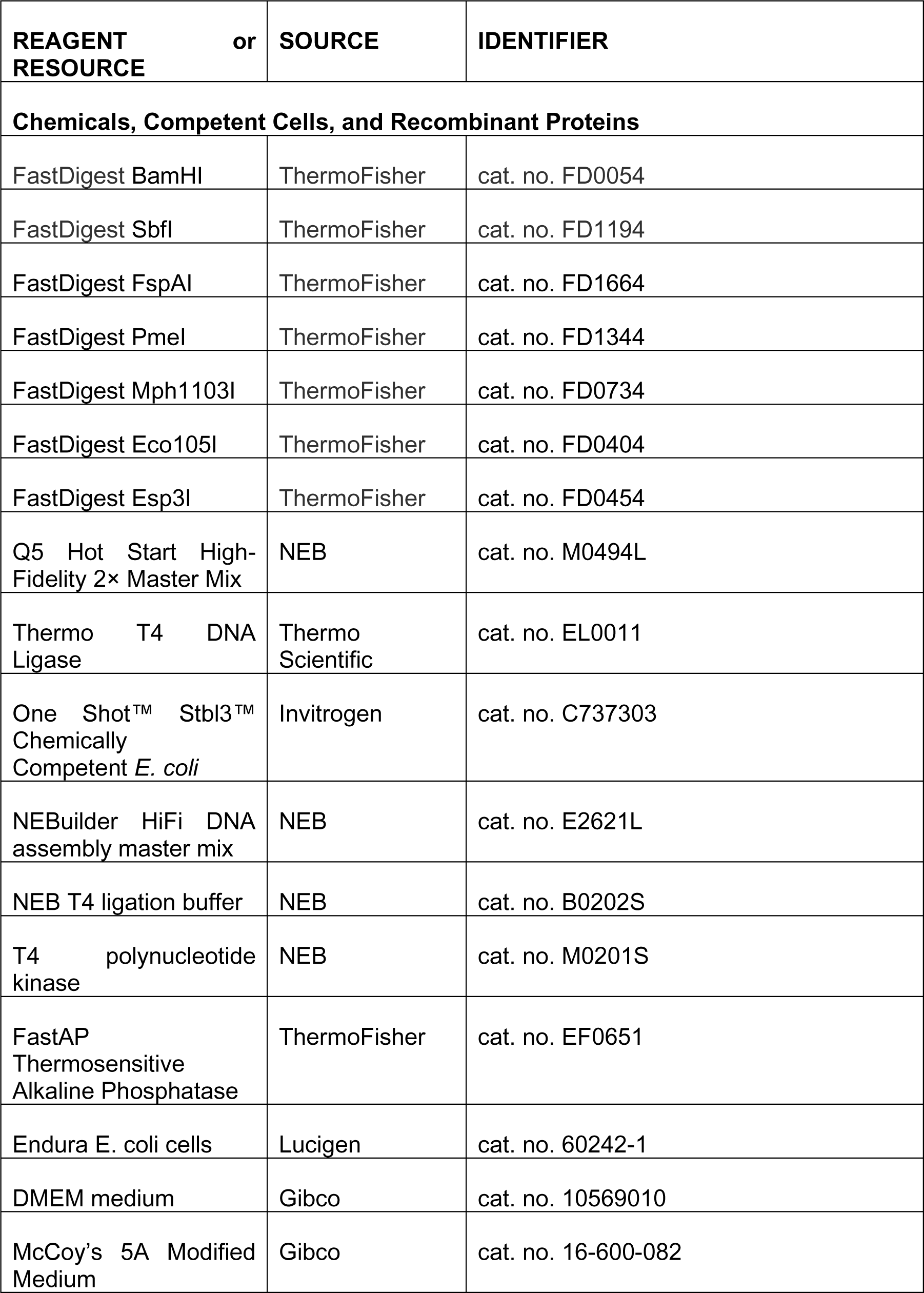
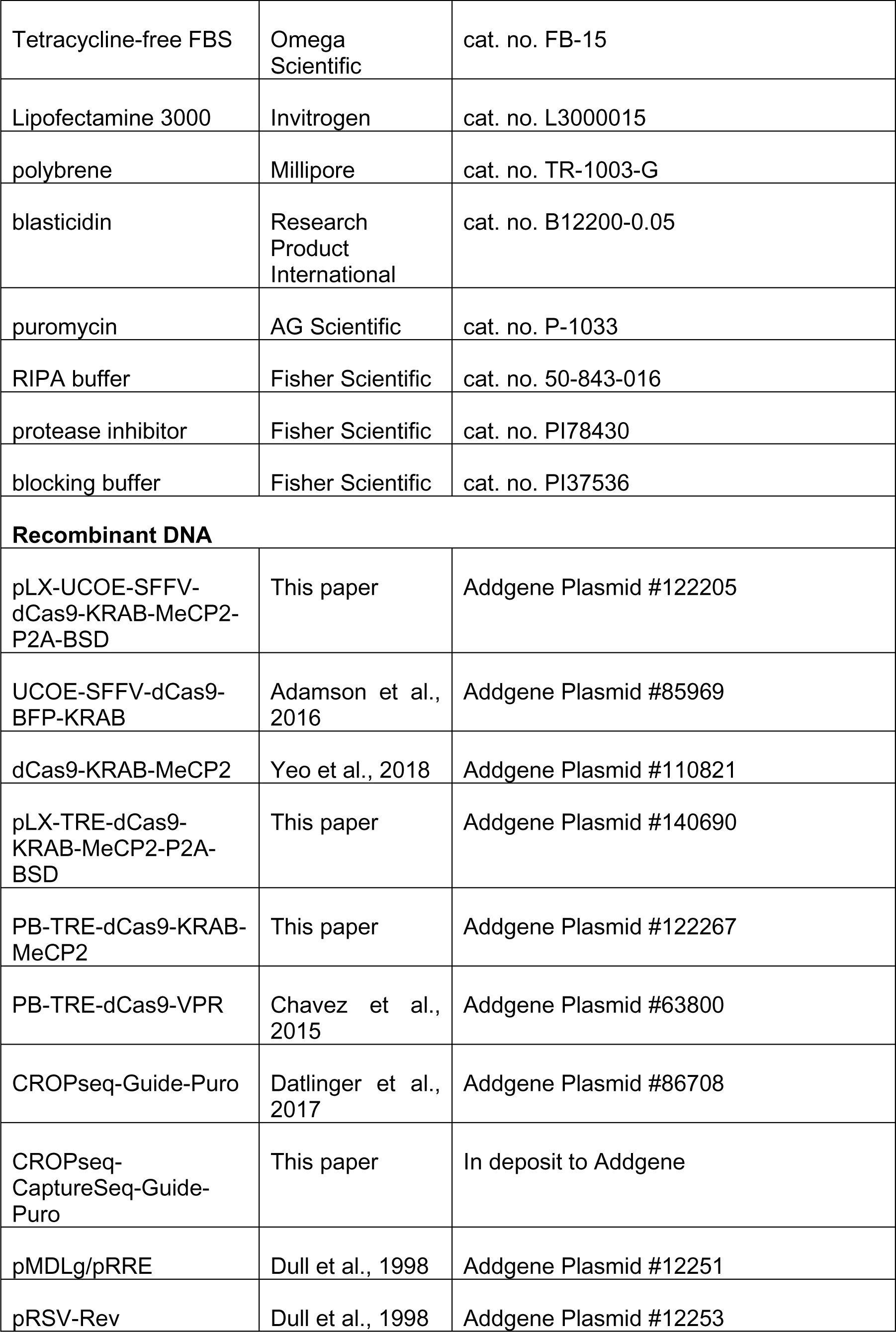
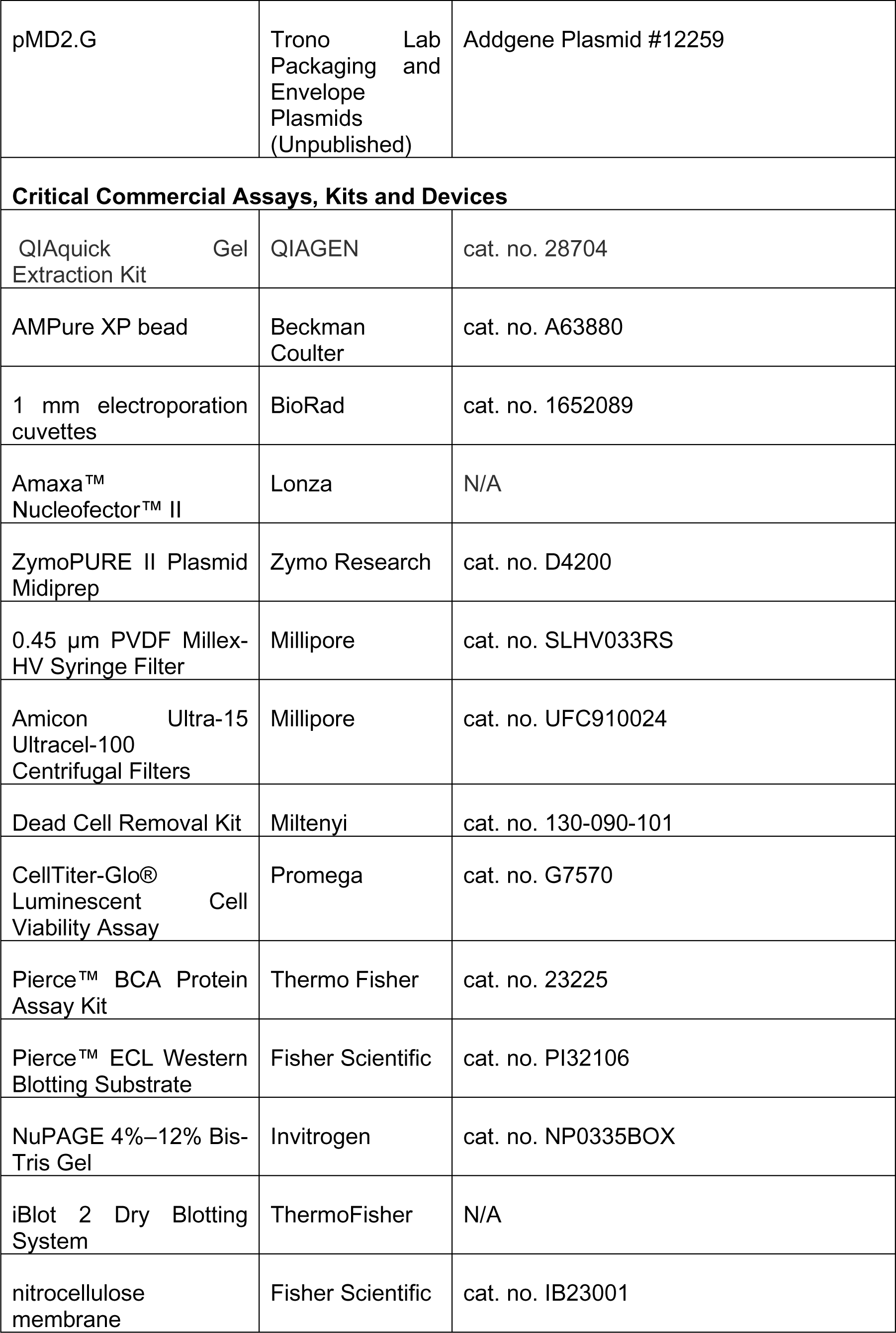
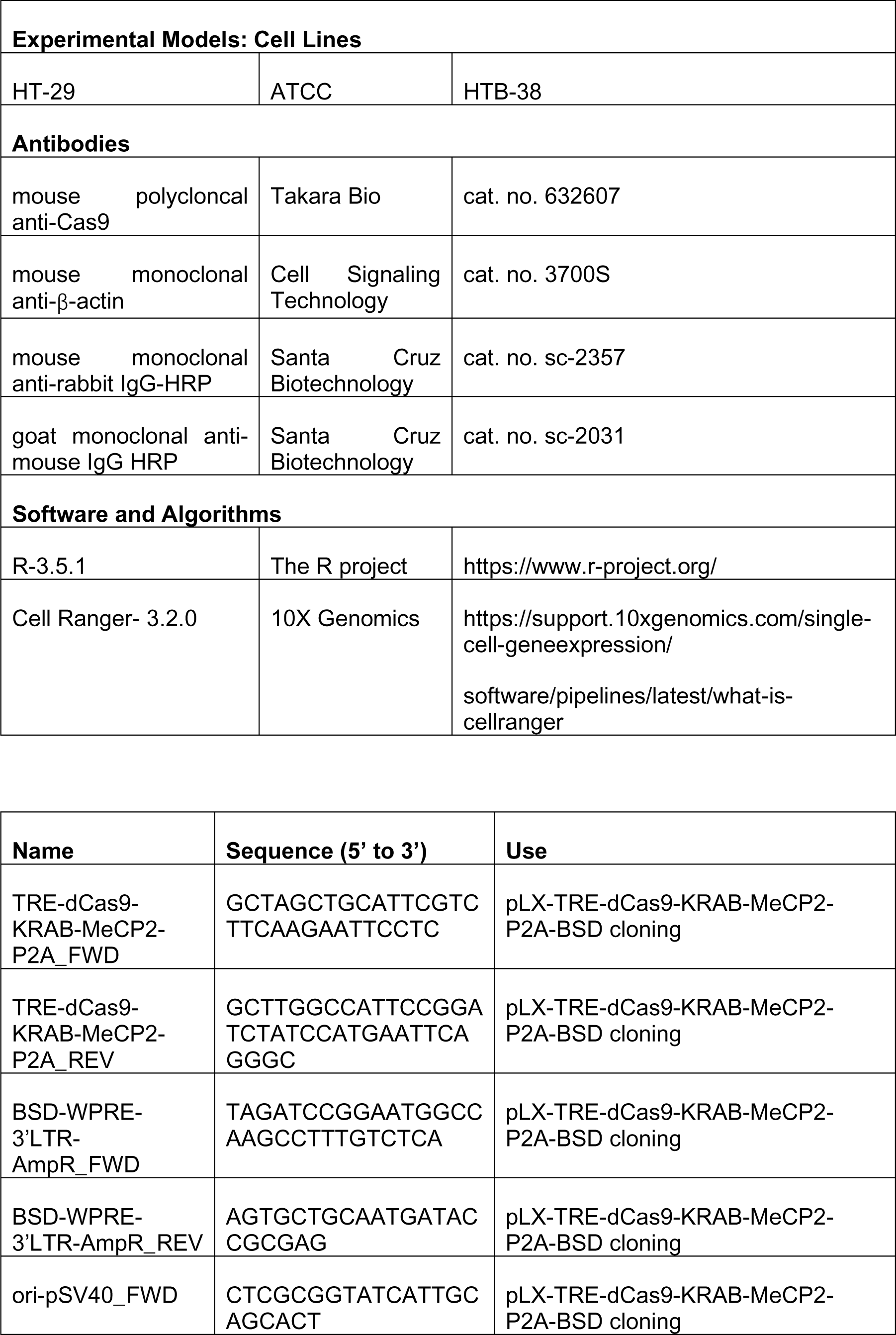
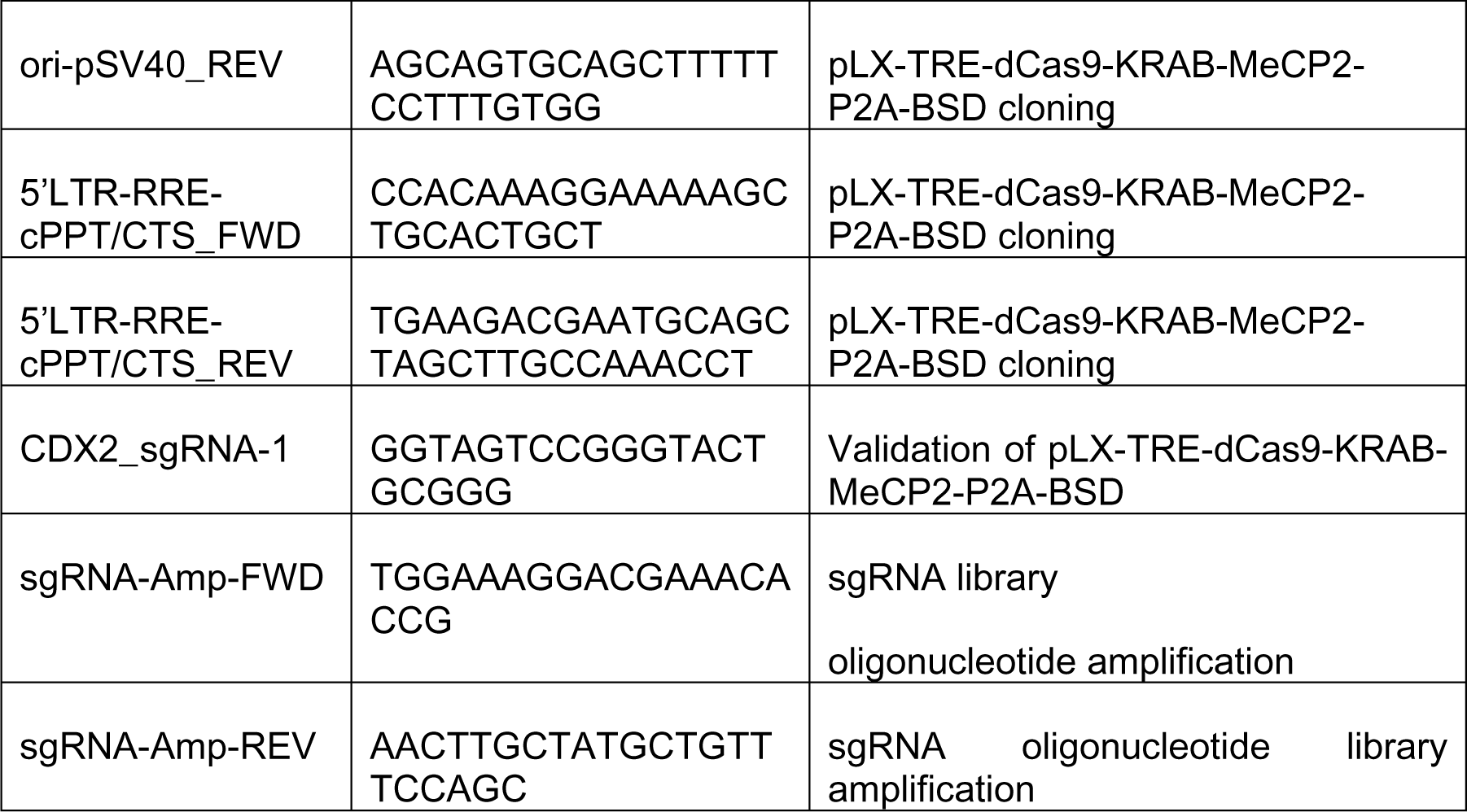

### Experimental procedures

#### Cloning and validation of the pLX-UCOE-SFFV-dCas9-KRAB-MeCP2-P2A-BSD plasmid

To clone the pLX-UCOE-SFFV-dCas9-KRAB-MeCP2-P2A-BSD plasmid, we prepared the vector backbone by incubating 5 ug of UCOE-SFFV-dCas9-BFP-KRAB (Addgene #85969) for 2 h at 37 °C with 2 ul of FastDigest BamHI (ThermoFisher cat. no. FD0054) and 2 ul of FastDigest SbfI (ThermoFisher cat. no. FD1194) in a total volume of 50 μl 1× ThermoFisher Green FastDigest Buffer. The digested product was run on a 0.8% agarose gel. The 13,792 bp fragment was cut under UV-free blue light and purified using the QIAquick Gel Extraction Kit (QIAGEN cat. no. 28704). The KRAB-MeCP2-P2A-BSD cassette was synthesized by Integrated DNA Technology as 1,634 bp dsDNA fragments with a BamHI restriction site on 5’ end and a SbfI restriction site on 3’ end. The synthesized product was digested by incubation for 1 h at 37 °C with 1 ul of FastDigest BamHI (ThermoFisher cat. no. FD0054) and 1 ul of FastDigest SbfI (ThermoFisher cat. no. FD1194) in a total volume of 20 μl 1× ThermoFisher Green FastDigest Buffer. The digested product was run on a 1.6% agarose gel. The 1,606 bp fragment was cut under UV-free blue light and purified using the QIAquick Gel Extraction Kit (QIAGEN cat. no. 28704). The vector backbone and KRAB-MeCP2-P2A-BSD cassette were ligated by the following T4 ligase reaction: 100 ng of vector backbone, 50 ng of KRAB-MeCP2-P2A-BSD cassette, 2 μl 10× Thermo Scientific T4 DNA Ligase Buffer, 2 μl 50% PEG 4000 solution, 1 μl T4 DNA Ligase (Thermo Scientific cat. no. EL0011) and water up to 20 μl, incubated at room temperature for 1 h. 5 μl of the ligation product were chemically transformed into Invitrogen™ One Shot™ Stbl3™ Chemically Competent E. coli (Invitrogen cat. no. C737303) following the manufacturer’s high-efficiency protocol. We screened for correctly assembled clones by colony PCR and further validated the assembly by restriction digestion with FspI, SbfI, AgeI and BamHI as well as Sanger sequencing.

#### Cloning and validation of the pLX-TRE-dCas9-KRAB-MeCP2-P2A-BSD plasmid

To clone the pLX-TRE-dCas9-KRAB-MeCP2-P2A-BSD plasmid, we first prepare the vector backbone by incubating 5 ug of PB-TRE-dCas9-VPR (Addgene #63800) for 2 h at 37°C with 2 ul of FastDigest FspAI (ThermoFisher cat. no. FD1664) and 2 ul of FastDigest PmeI (ThermoFisher cat. no. FD1344) in a total volume of 50 μl 1× ThermoFisher Green FastDigest Buffer. The digested product was run on a 0.8% agarose gel. The 10,992 bp was cut under UV-free blue light and purified using the QIAquick Gel Extraction Kit (QIAGEN cat. no. 28704). The truncated-dCas9-KRAB-MeCP2 cassette was synthesized by Integrated DNA Technology as 2,450 bp dsDNA fragments with a FspAI restriction site on 5’ end and a PmeI restriction site on 3’ end. The synthesized product was digested by incubation for 1 h at 37 °C with 1 ul of FastDigest FspAI (ThermoFisher cat. no. FD0054) and 1 ul of FastDigest PmeI (ThermoFisher cat. no. FD1194) in a total volume of 20 μl 1× ThermoFisher Green FastDigest Buffer. The digested product was run on a 1.6% agarose gel. The 2,432 bp fragment was cut under UV-free blue light and purified using the QIAquick Gel Extraction Kit (QIAGEN cat. no. 28704). The vector backbone and truncated-dCas9-KRAB-MeCP2 cassette were ligated by the following T4 ligase reaction: 100 ng of vector backbone, 100 ng of truncated-dCas9-KRAB-MeCP2 cassette, 2 μl 10× Thermo Scientific T4 DNA Ligase Buffer, 2 μl 50% PEG 4000 solution, 1 μl T4 DNA Ligase (Thermo Scientific cat. no. EL0011) and water up to 20 μl, incubated at room temperature for 1 h. 5 μl of the ligation product were chemically transformed into Invitrogen™ One Shot™ Stbl3™ Chemically Competent E. coli (Invitrogen cat. no. C737303) following the manufacturer’s high-efficiency protocol. We screened for correctly assembled clones by colony PCR and further validated the assembly by restriction digestion with SalI, FspAI and PmeI as well as Sanger sequencing.

To clone the pLX-TRE-dCas9-KRAB-MeCP2-P2A-BSD plasmid, the TRE-dCas9-KRAB-MeCP2-P2A cassette (7,500 bp) was amplified from PB-TRE-dCas9-KRAB-MeCP2 and purified using protocol described above. The pLX vector backbone was amplified as 3 fragments (BSD-WPRE-3’LTR-AmpR cassette as 3,066 bp fragment, ori-pSV40 cassette as 2,942 bp fragment, 5’LTR-RRE-cPPT/CTS cassette as 3,543 bp fragment) from pLX-UCOE-SFFV-dCas9-KRAB-MeCP2-P2A-BSD and purified using the protocol described above. Primers for amplification were designed to contain >15bp homology arm on each side with adjacent fragments. The 4 fragments were assembled using Gibson’s isothermal assembly: 500ng of total amplified fragments were combined with 10μl of NEBuilder HiFi DNA assembly master mix (NEB cat. no. E2621L) and water to 20μl. After 1 h of incubation at 50 °C, reactions were purified by AMPure XP bead clean-up (Beckman Coulter cat. no. A63880), and 5μl of the ligation product were chemically transformed into Invitrogen™ One Shot™ Stbl3™ Chemically Competent E. coli (Invitrogen cat. no. C737303) following the manufacturer’s high-efficiency protocol. We screened for correctly assembled clones by colony PCR and further validated the assembly by restriction digestion with AdeI, FspAI and PmeI as well as Sanger sequencing.

#### Cloning and validation of the CROPseq-CaptureSeq-Guide-Puro plasmid

To clone the CROPseq-CaptureSeq-Guide-Puro plasmid, we prepared the vector backbone by incubating 5 ug of CROPseq-Guide-Puro (Addgene #86708) for 2 h at 37 °C with 2 ul of FastDigest Mph1103I (ThermoFisher cat. no. FD0734) and 2 ul of FastDigest Eco105I (ThermoFisher cat. no. FD0404) in a total volume of 50 μl 1× ThermoFisher Green FastDigest Buffer. The digested product was run on a 0.8% agarose gel. The 9,766 bp fragment was cut under UV-free blue light and purified using the QIAquick Gel Extraction Kit (QIAGEN cat. no. 28704). The trcrRNA-CaptureSeq-5’LTR(truncated) cassette was synthesized by Integrated DNA Technology as 520 bp dsDNA fragments with 20 bp of homology on each end of the digested vector backbone, respectively.

The vector backbone and trcrRNA-CaptureSeq-5’LTR(truncated) cassette were assembled using Gibson’s isothermal assembly: 100ng of vector backbone and 20ng of trcrRNA-CaptureSeq-5’LTR(truncated) cassette were combined with 10 μl of NEBuilder HiFi DNA assembly master mix (NEB cat. no. E2621L) and water to 20 μl. After 1 h of incubation at 50 °C, reactions were purified by AMPure XP bead clean-up (Beckman Coulter cat. no. A63880), and 5 μl of the ligation product were chemically transformed into Invitrogen™ One Shot™ Stbl3™ Chemically Competent E. coli (Invitrogen cat. no. C737303) following the manufacturer’s high-efficiency protocol. We screened for correctly assembled clones by colony PCR and further validated the assembly by restriction digestion with Mph1103I, AdeI and BsmbI as well as Sanger sequencing.

### Cloning of individual sgRNAs into the CROPseq-CaptureSeq-Guide-Puro plasmid

sgRNA cassettes were annealed from two oligonucleotides (top: 5’-CACCG(N)_19-20_-3’ bottom: 5’-AAAC(N)_19-20_-3’) by combining 1 μl of each 100 μM oligonucleotide with 1 μl of 10× T4 ligation buffer (NEB cat. no. B0202S), 6.5 μl of water, and 0.5 μl of T4 polynucleotide kinase (NEB cat. no. M0201S), incubating as follows: 37 °C for 30 min (oligonucleotide phosphorylation), 95 °C for 5 min, then ramping from 90 °C to 25 °C at 5 °C/min. Vector backbone was prepared by digesting and dephosphorylating 5 μg of CROPseq-CaptureSeq-Guide-Puro with 5 μl of FastDigest Esp3I (ThermoFisher cat. no. FD0454) and 2 μl of FastAP Thermosensitive Alkaline Phosphatase (ThermoFisher cat. no. EF0651) in a total volume of 50 μl 1× ThermoFisher Green FastDigest Buffer, incubating for 1 h at 37 °C. The digested product was run on a 1.6% agarose gel. The 8,361 bp fragment was cut under UV-free blue light and purified using the QIAquick Gel Extraction Kit (QIAGEN cat. no. 28704).

Ligation reactions were set up as follows: 60 μg of CROPseq-CaptureSeq-Guide-Puro backbone, 1 μl gRNA cassette (diluted 1:200 in water), 2 μl 10× Thermo Scientific T4 DNA Ligase Buffer, 2 μl 50% PEG 4000 solution, 1 μl T4 DNA Ligase (Thermo Scientific cat. no. EL0011) and water up to 20 μl, incubated at room temperature for 1 h. The ligation reaction was chemically transformed into Invitrogen™ One Shot™ Stbl3™ Chemically Competent E. coli (Invitrogen cat. no. C737303) following the manufacturer’s high-efficiency protocol.

### Cloning of pooled sgRNA libraries into the CROPseq-CaptureSeq-Guide-Puro plasmid

Vector backbone was prepared by digesting 5 μg of CROPseq-CaptureSeq-Guide-Puro with 5 μl of FastDigest Esp3I (Thermo Scientific cat. no. FD0454) in a total volume of 50 μl 1× Thermo Scientific FastDigest Green Buffer, incubating for 1 h at 37 °C. The digested product was run on a 1.6% agarose gel. The 8,361 bp fragment was cut under UV-free blue light and purified using the QIAquick Gel Extraction Kit (QIAGEN cat. no. 28704).

sgRNA insert fragments were synthesized by TWIST Bioscience as 74 nt oligonucleotides with 18 and 35 nt of homology to the hU6 promoter and guide RNA scaffold, respectively. Oligonucleotides were diluted to 1 ng/ul and pooled in equal amounts. The oligonucleotide pool was further amplified by mixing 12.5 μl KAPA HiFi HotStart 2× ReadyMix (KAPA Biosystems cat. no. KK2601), 0.75 μl of 10 μM forward primer, 0.75 μl of 10 μM reverse primer, 2 ng oligonucleotide pool and water up to 25 μl and incubating as follows: 95 °C for 3 min, 9× (98 °C for 20 s, 63 °C for 15 s, 72 °C for 15 s), 72 °C for 1 min, hold at 4 °C. PCR product was run on a 2% agarose gel. The 74 bp fragment was cut under UV-free blue light and purified using the QIAquick Gel Extraction Kit (QIAGEN cat. no. 28704).

sgRNA libraries were cloned by Gibson’s isothermal assembly: 22 fmoles (113.7 ng) of CROPseq-CaptureSeq-Guide-Puro backbone and 400 fmoles (18.31ng) of amplified dsDNA oligonucleotides were combined with 10 μl of NEBuilder HiFi DNA assembly master mix (NEB cat. no. E2621L) and water to 20 μl. After 1 h of incubation at 50 °C, reactions were purified by AMPure XP bead clean-up (Beckman Coulter cat. no. A63880), and 10 μl of the reaction was electroporated into 50 μl of Lucigen Endura E. coli cells (Lucigen cat. no. 60242-1) using prechilled 1 mm electroporation cuvettes (BioRad cat. no. 1652089) in a Lonza Amaxa™ Nucleofector™ II device set to Bacteria Program 4. Within seconds after the pulse, 1 ml of 37 °C Lucigen Recovery Medium was added and the bacteria were grown in a round-bottom tube for 1 h at 37 °C while shaking at 200 r.p.m. Then, 1 ml of the bacterial culture was plated on a 25 × 25 cm bioassay plate containing LB medium (Miller) with 100 μg/ml carbenicillin. Plates were incubated at 30 °C for 20 h, then LB medium was added and bacteria colonies were scraped off the plate. Bacterial cells were pelleted by 30 min of centrifugation at 3,000 RCF at 4 °C, and plasmid DNA was extracted with multiple ZymoPURE II Plasmid Midiprep columns (Zymo Research cat. no. D4200). Library coverage was estimated by counting the number of bacterial colonies on a 1:1,000 dilution plate and a 1:10,000 dilution plate. All the sgRNA libraries were cloned with at least 500× coverage.

### Lentivirus production for single sgRNAs and pooled CROP-seq screens

HEK293T cells were plated into 100mm dishes at 2 million cells per dish in 12 ml of culture medium (DMEM (Gibco cat. no. 10569010), 10% Tetracycline-free FBS (Omega Scientific cat. no. FB-15), no antibiotics) and grown to reach 50% to 70% confluence. 6ml of culture medium was removed from the dishes and the cells were transfected with lipofectamine 3000 (Invitrogen cat. no. L3000015) using either 10.2 μg of CROPseq-CaptureSeq-Guide-Puro (containing single sgRNAs or libraries) or dCas9 expressing plasmid, and 4.5 μg each of the three packaging plasmids pMDLg/pRRE (Addgene #12251), pRSV-Rev (Addgene #12253), and pMD2.G (Addgene #12259). The medium was exchanged for fresh culture medium 16 h after the transfection. The supernatant containing viral particles was harvested at 30 h and passed through a 0.45 µm PVDF Millex-HV Syringe Filter (Millipore cat. no. SLHV033RS) to remove cells and debris. Viral particles were further concentrated by 40min centrifugation at 4,000 r.p.m. using Amicon Ultra-15 Ultracel-100 Centrifugal Filters (Millipore cat. no. UFC910024). The concentrated virus was aliquoted and stored at −80 °C.

### Production of constitutive/inducible dCas9 expressing cell lines

For adherent cell lines HT-29, cells were seeded into 100mm dishes at 5 million cells per dish in 12 ml of complete culture medium (McCoy’s 5A Modified Medium for HT-29 (Gibco cat. no. 16-600-082) with 10% Tetracycline-free FBS (Omega Scientific cat. no. FB-15), no antibiotics). The cells were then transduced with 8 μg/ml polybrene (Millipore cat. no. TR-1003-G) and the proper amount of lentivirus containing dCas9 vector to ensure Multiplicity of Infection (MOI) equal to 1. An extra dish served as the untransduced control. After addition of the virus, cells were incubated overnight at 37 °C, 5% CO2. At 24 h post-transduction, selection with 20 μg/ml blasticidin (Research Product International cat. no. B12200-0.05) began. After 5 days of blasticidin selection, live cells were trypsinized and seeded into 100mm dishes at 2 million cells per dish in fresh complete culture medium containing 10 μg/ml blasticidin to allow for cell number amplification while renewing the medium (containing blasticidin) every 3 days.

### Lentiviral transduction with sgRNA libraries or single sgRNAs

For adherent cell lines HT-29, cells were seeded into 100mm dishes at 5 million cells per dish in 12 ml of complete culture medium (McCoy’s 5A Modified Medium for HT-29 (Gibco cat. no. 16-600-082) with 10% Tetracycline-free FBS (Omega Scientific cat. no. FB-15), no antibiotics) containing 10 μg/ml blasticidin (Research Product International cat. no. B12200-0.05) to maintain selection for dCas9 expression. The cells were then transduced with 8 μg/ml polybrene (Millipore cat. no. TR-1003-G) and the proper amount of lentivirus containing sgRNA libraries to ensure Multiplicity of Infection (MOI) equal to 0.3. An extra dish served as the untransduced control. After addition of the virus, cells were incubated overnight at 37 °C, 5% CO2. At 24 h post-transduction, selection with 8 μg/ml puromycin (AG Scientific cat. no. P-1033) began. After 3 days of puromycin selection, live cells were trypsinized and seeded into 100mm dishes at 2 million cells per dish in fresh complete culture medium containing blasticidin and 4 μg/ml puromycin. After overnight incubation at 37 °C, 5% CO2 to allow for reattaching, the cells are ready for doxycycline induction.

### Lentivirus titration for single sgRNAs

Cells were seeded into 24-well plates at 50,000 cells per well in 500 μl of proper culture medium and grown overnight to reach 30% to 50% confluence. The next day, medium was exchanged for 450 μl per well of fresh culture medium containing 8 μg/ml polybrene (Millipore cat. no. TR-1003-G), which was also used to dilute the viral stock. Lentivirus aliquots were thawed from storage at −80 °C and titrated in a 1:2 dilution series ranging over six wells (1:2 to 1:32). Each dilution was tested in duplicate by adding 50 μl per well. At least two wells per dilution served as un-selected controls and at least two wells per plate served as untransduced controls. 24 h after the transduction, the medium was exchanged for 500 μl per well of selection medium prepared as described above, every 2–3 days. The blasticidin selection took 5 days and the puromycin selection took 3 days for HT-29. As soon as the selection was completed and all cells in the untransduced controls had died, cell viability was measured by CellTiter-Glo® Luminescent Cell Viability Assay (Promega cat. no. G7570). The virus titer (transducing units per ml) was calculated as follows: (initial number of cells × percentage of viable cells × dilution factor) × (1,000 μl /50 μl).

### Quantification of dCas9 by Western Blot

To evaluate inducible dCas9 expression, HT-29 cells were transduced with pLX-TRE-dCas9-KRAB-MeCP2-P2A-BSD as described above. Transduced HT-29 were then plated in RPMI medium (Gibco cat. no. 21875-034) on 6-well plates at 0.5 million cells per well. Cells were recovered overnight and doxycycline (Fisher Scientific cat. no. BP26535) was added to culture medium at 1 μg/ml. At 6, 9, 12 and 24 h post doxycycline induction, respectively, cells were lysed in 300 μl RIPA buffer (Fisher Scientific cat. no. 50-843-016) with protease inhibitor (Fisher Scientific cat. no. PI78430). Protein concentration was measured by Pierce™ BCA Protein Assay Kit (Thermo Fisher cat. no. 23225). 20 ug of total protein from each lysate was loaded and separated on a NuPAGE 4%–12% Bis-Tris Gel (Invitrogen cat. no. NP0335BOX). Subsequently, the protein was transferred onto a nitrocellulose membrane (Fisher Scientific cat. no. IB23001) by iBlot 2 Dry Blotting System (ThermoFisher), which was then blocked by blocking buffer (Fisher Scientific cat. no. PI37536) for 1 h at room temperature. The blocked membrane was incubated overnight at 4 ℃ in primary antibodies: mouse polyclonal anti-Cas9 (Takara Bio, cat. no. 632607), mouse monoclonal anti-β-actin (8H10D10) (Cell Signaling Technology, cat. no. 3700S). After incubation, the membrane was washed 3 times with PBS-T buffer and then incubated at room temperature for 1 h with secondary antibodies: mouse monoclonal anti-rabbit IgG-HRP (Santa Cruz Biotechnology, cat. no. sc-2357), goat monoclonal anti-mouse IgG HRP (Santa Cruz Biotechnology, cat. no. sc-2031). The membrane was then washed 3 times with PBS-T buffer, activated by Pierce™ ECL Western Blotting Substrate (Fisher Scientific cat. no. PI32106), and imaged by X-Ray film. The film was then scanned and digital images were processed using ImageJ.

### Computational procedures

#### Preprocessing of single-cell sequencing data

Single-cell sequencing data were aligned using CellRanger v3.0.2 to the human reference genome assembly (Ensembl GRCh38 release) with default parameters. CellRanger pipeline uses STAR for alignment. Briefly, reads aligned to exons are tagged with their respective gene names (annotated by Ensembl Gene ID). Then counts of unique molecular identifier (UMI)-deduplicated reads per gene within the same cell were counted to build a gene expression matrix comprising all cells with UMI counts. For downstream analysis, low expression genes–defined by being detected in less than 10% cells–are discarded. We found this step is critical for controlling noise. Furthermore, as quality control at single-cell level, cells with either low transcriptome complexity (<5000 UMI counts) or multiple sgRNAs were discarded. Cells with high mitochondrial reads fraction (>15%) were excluded in differential expression analysis and further downstream analyses.

### Assignment of sgRNA to single-cell transcriptome

In the setting of this project, we enabled the detection of two types of sgRNA transcripts by incorporating Capture Sequence (Replogle, Norman et al. 2020) into the CROP-Seq sgRNA expressing vector CROP-Seq-GuidePuro (Datlinger, Rendeiro et al. 2017). Polyadenylated sgRNA transcripts synthesized by RNA polymerase II were captured by oligo-dT primers and subsequently sequenced as part of whole-transcriptome library. Non-polyadenylated sgRNA transcripts synthesized by RNA polymerase III were captured by CaptureSeq primers and were separately prepared as Feature Barcode libraries following 10X Genomics protocol (User Guide for Chromium Single Cell 3ʹ Reagent Kits v3). Feature Barcode libraries were separately indexed and sequenced as spike-ins alongside the whole-transcriptome single-cell RNA-seq libraries. Final UMI and cell barcode assignments were made for each Feature Barcode read by alignment with CellRanger v3.0.2, as was done for the whole transcriptome libraries.

To computationally detect both types of sgRNA transcripts, we adapted and modified the sgRNA identification approach described in (Hill, McFaline-Figueroa et al. 2018). We allowed a maximal Hamming distance of 1 between the protospacer sequences extracted from the sequencing reads and the input sgRNA library. To account for background sgRNA reads resulting from low level of sgRNA transcripts released from lysed cells, we adopted a 2-Gaussian Mixture Model strategy described in CellRanger’s “CRISPR Algorithm” (https://support.10xgenomics.com/single-cell-gene-expression/software/pipelines/latest/algorithms/crispr) and in (Replogle, Norman et al. 2020). We then made sgRNA assignments separately for whole-transcriptome library and Feature Barcode library. For each library, a cell was considered to express only one sgRNA if the most abundant sgRNA was five times more abundant than the second most abundant sgRNA; they were considered to express multiple sgRNAs if this ratio was smaller than five. To consolidate the sgRNA assignment from the two libraries, cells were considered to express unique sgRNA if 1) the same unique sgRNA was detected in the two libraries or 2) only one sgRNA is detected in one library and no sgRNA was detected in the other. Cells were considered to express multiple sgRNAs if 1) multiple sgRNAs were detected in either library or 2) different sgRNAs are detected in the two libraries.

This dual sgRNA detection strategy helps more thorough detection of sgRNA transcripts in single cells and effective removal of multiplets. We performed TReK experiments using MOI=0.3 to ensure that most of the single cells express only one sgRNA. As expected, for the majority (roughly 60% across two timepoints) of unique-sgRNA cells, the same sgRNA was identified from the two sgRNA-containing libraries. Our data also shows that while two libraries generated highly consistent sgRNA-cell association maps, each can detect sgRNAs the other strategy cannot detect in a small subset of cells (Figures S3E-S3G).

We observed that cells expressing only one sgRNA are enriched in the 10,000-50,000 UMI count range (Figures S3C and S3D). On the lower UMI count side, the majority of cells do not have any sgRNA detected due to lower transcriptome complexity. On the higher UMI count side, a significant proportion of cells have multiple sgRNAs detected, indicating that these transcriptomes likely resulted from droplets with multiple single cells, considering that the proportion of cells expressing multiple sgRNAs are theoretically minimal with low MOI transduction. The association between abnormally high transcriptome complexity and the detection of multiple sgRNAs indicates that with the dual sgRNA detection strategy, we were able to retrospectively remove potential doublets and triplets—a source of significant noise in single-cell sequencing--by simply removing cells with multiple sgRNAs.

### Determination of on-target knockdown efficacy

To measure the extent to which each sgRNA reduces the expression of its target gene (knockdown efficacy), we compared cells expressing the corresponding sgRNA to cells expressing scrambled sgRNAs (sgCtrls). For each sgRNA, we estimated its target gene expression level by aggregating all cells that express the corresponding sgRNA. We also estimated the unperturbed expression level by aggregating the same number of randomly sampled cells that express sgCtrls. The sampling was done 100 times and the median value was taken. The knockdown efficacy of the sgRNA was then calculated as the fraction of target gene expression (normalized as count per million, or CPM) that was reduced due to the presence of the corresponding sgRNA.

Broadly, we observed acceptable silencing for most genes, with 1,877/3,016 sgRNAs (62%) achieving ≥ 50% knockdown and 1,275/3,016 sgRNAs (42%) achieving at least 80% knockdown at 96h (Figures S4A and S4B; Table S7; Methods). Since 3 sgRNAs per TR were used, most TRs were silenced at different silencing efficacies (Figures S4C and S4D). This turned out to be a useful feature, for instance, to dissect compensatory, dosage-dependent effects (Figure 4). When comparing silencing efficacy between timepoints, we generally observed a stronger knockdown at T2 = 96h vs. T1 = 24h (Figures S4A and Figure S4B). While this difference may be partly due to differences in the strengths of the constitutive and inducible promoters, we believe that much of the reduced efficacy at T1 is due to a combination of two factors: First, the dCas9-KRAB-MeCP2 abundance necessary for effective knockdown may vary from gene to gene. Second, mRNA degradation kinetics and the size of the extant mRNA pool can vary widely across genes.

### Differential expression analysis

To perform differential expression analysis for each sgRNA perturbation, a pseudo-bulk sample was generated by aggregating the transcriptomes of all the single cells expressing that particular sgRNA. A pseudo-bulk gene expression signature was obtained by comparing the expression level of each gene against 200 reference pseudo-bulk samples as 1) z-scores generated by viperSignature function from viper R package and 2) log fold change. The reference pseudo-bulk samples were obtained by aggregating the transcriptomes of randomly sampled cells expressing other sgRNAs. To control for the variation resulted from cell number, each reference pseudo-bulk sample was aggregated from the same number of cells as the “tested” pseudo-bulk sample.

To perform differential expression analysis for each perturbed TR (as shown in Figures 2C, 3, S5G, and S5H), we first grouped cells expressing any of the 3 sgRNAs targeting that particular TR. These cells were then sorted by the normalized expression level of the perturbed TR. The top 50% of the cells are then selected to represent the TR-perturbed cell population. (If two cells have the same level of TR expression, the cell expressing more efficient sgRNA was selected. If two cells express the same sgRNA, the cell with a higher total UMI count was selected.) The differential expression analysis for the particular TR was then conducted in the same way as described in sgRNA-wise differential expression analysis.

### Assessing the reproducibility of TReK differential expression

An unbiased approach to assess gene knockdown reproducibility is to compare different technologies. Searching the published literature, we found that RNA-sequencing of HT-29 cells following CRISPRi-mediated TCF7L2 silencing had been performed to investigate this gene’s role in colorectal cancer invasion and migration (Wenzel, Rose et al. 2020). Impressively, even when pooling only 17 or 21 cells (at T1 and T2, respectively), the overlap in terms of differentially expressed genes between our data and Wenzel et al. data is highly significant (empirical p < 0.001) (Figures S5G and S5H).

We were interested in the factors that affect the correlation between the differentially expressed gene signatures induced by different sgRNAs targeting the same gene. Not surprisingly, our analysis confirmed that the TRs with the most correlated gene signatures (Figures S5A and S5B) tended to be those with multiple high-efficiency sgRNAs (Figure S5C). Additionally, TRs with multiple, high-efficiency sgRNAs affected a greater fraction of downstream transcriptional targets (Figure S5D) and were characterized by slightly higher coverage (cells per sgRNA) (Figure S5E).

We were also interested in the effect of protein turnover on differential gene expression at early timepoints. For instance, one would expect that proteins with low turnover should affect a greater number of downstream target genes at T2 vs. T1 compared to proteins with high turnover. To investigate, we collected published data on protein stability as measured by mass-spectrometry (Cambridge, Gnad et al. 2011). Although we could find protein stability data for only a relatively small fraction of the TRs silenced in our assays, we observed that more stable proteins tend to have a weaker differential expression at the earlier timepoint T1 compared to T2 and thus a higher intra-gene sgRNA correlation at T2 vs. T1 (Figure S5F).

### Gene Set Enrichment Analysis

One-set Gene Set Enrichment Analysis (GSEA) was implemented as described in (Subramanian, Tamayo et al. 2005). Briefly, the gene expression signature of interest was sorted and scanned by calculating an enrichment score (ES) starting with the most upregulated gene. If the encountered gene was present in the gene set, the running ES was increased. If the encountered gene was not present in the gene set, then the running ES was decreased. The final ES was determined by the maximum running ES and the leading edge subset was defined as the genes that were encountered before reaching maximum ES. The statistical significance of the ES was calculated by permuting differential expression-ordered genes 1,000 times, computing the ES to generate a null distribution, and comparing the unpermuted ES score to the null distribution of ES. Normalized enrichment score (NES) was calculated as the z-score transformed ES against the null distribution. An empirical p-value was calculated as the fraction of permuted ES that were more extreme than the actual ES.

For two-set GSEA (Figures S5G and S5H), the query gene set was divided into two subsets: a positive subset containing genes that were positively correlated with the gene set term, and a negative subset containing genes that were negatively correlated with the gene set term. The gene expression signature of interest was sorted and scanned as described in the one-set GSEA process. However, the ES for the positive and negative subset, respectively, were calculated separately and subsequently added together. The NES and empirical p-values were computed as described above.

### Determination of mode of regulation (Figure 4F, Table S6)

To determine the mode of regulation of a TR within a specific pathway, GSEA analysis was conducted on the gene expression signature of all 3 sgRNAs targeting the particular TR. If the NES of at least two sgRNAs were statistically significant (*p* < 0.05) and have the same sign, the NES of the third sgRNA was taken into account. If the third sgRNA perturbation showed statistically significant NES with the same sign or 2) showed no statistically significant NES, the mode of regulation was determined by the first two sgRNAs. If the third sgRNA perturbation showed statistically significant NES with opposite sign, the mode of regulation was determined as “mixed”.

### ARACNe network reconstruction

For each sgRNA, a pseudo-bulk sample was generated by aggregating transcriptomes of 15 single cells expressing that particular sgRNA. To maximize the transcriptome coverage, we included sgRNAs covered by more than 15 cells. This procedure generated 1,447 pseudo-bulk expression profiles for 14,773 transcriptionally informative genes (defined as being expressed by >10% samples) from T1 dataset and 1,310 pseudo-bulk expression profiles for 18,426 transcriptionally informative genes from T2 dataset. These expression profiles were used to generate ARACNe networks (“purely computational networks”) for transcription factors (TFs), coTFs and signaling proteins as previously described in (Basso, Margolin et al. 2005).

### Hurdle model-based generalized linear model

Hurdle model was firstly described in (Cragg 1971) and applied in the context of single-cell RNA-seq in (Finak, McDavid et al. 2015) as MAST (Model-based Analysis of Single-cell Transcriptomics) approach. Here we modified MAST to be compatible with the single-cell CRISPRi perturbation setting. Normalized gene expression (count per million or CPM) was modeled as a generalized regression model (GLM) with two branches, one in binary space and the other in continuous space. For a specific gene, the probability of detecting non-zero CPM in a cell was modeled by binomial distribution, whereas the exact CPM|CPM>0 was modeled as Poisson distribution.

Besides the intercept unit in the GLM, we included two covariates in the regression. The first covariate represents the expression level of the TR of interest. When its expression level was unavailable due to low expression, protein activity inferred by VIPER was used to estimate the abundance of the TR. Secondly, as several publications found that the consideration of “cellular detection rate” (CDR), defined as the proportion of genes detected in a single cell, significantly improved the differential expression analysis in single-cell sequencing context (Finak, McDavid et al. 2015, Dixit, Parnas et al. 2016, Soneson and Robinson 2018), we included CDR as the second covariate.

Regression coefficients were estimated and regularized as described in (Finak, McDavid et al. 2015). For the binary model, coefficients were regularized using informative priors (Gelman et al., 2008) from which information from the ARACNe network can be incorporated. In practice, we assigned a 4x larger prior variance to any TR-target associations that were identified by the ARACNe network. For each TR, only a subset of the transcriptome was considered as candidate target genes. For the “integrated network”, the candidate target genes came from two sources: 1) target genes identified by ARACNe network and 2) the top 200 most differentially expressed genes after the TR perturbation. For the “purely experimental network”, the top 200 most differentially expressed genes after the TR perturbation were considered as candidate target genes.

### Determination of regulon parameters

We define the “regulon” of a specific TR as all the transcriptional target genes of that TR. To generate regulons that can be used to infer protein activity of a TR, two sets of parameters have to be determined: tfmode and likelihood. As described in (Alvarez, Shen et al. 2016), the tfmode parameter represents the direction as well as strength of the TR-target regulation. The likelihood parameter denotes the statistical confidence of the TR-target regulation. For the “purely computational” ARACNe network, tfmode was calculated as the Pearson correlation of the corresponding regulator and target gene expression, and likelihood was calculated as scaled pairwise mutual information. For “purely experimental” networks and integrated networks, the linear model coefficient statistics were used to generate calculate the regulon parameters. We first determined the statistical significance of each coefficient assuming a Gaussian distribution. For each TR-target association in the regulon, the use of the continuous versus binary model was determined by greater coefficient significance. The tfmode parameter was then determined as the sign (1 or -1) of the coefficient. The likelihood parameter was determined as the significance (1 - p-value) of the corresponding coefficient.

### 5-fold cross-validation and cross-timepoint validation

Within the dataset of each timepoint, 5-fold cross-validation was used to assess the ability of a specific regulon to infer the protein activities of the corresponding TR. For each TR, a single-cell dataset was firstly assembled by including all the perturbed cells and cells expressing a scrambled sgRNA. The dataset was subsequently divided into 5 equally-sized sub-datasets. The regulon was reconstructed from 4 sub-datasets (training sets) and validated on the single left-out sub-dataset (test set). The protein activities of the TR in each perturbed cell were then calculated by VIPER based on computational network, experimental network or integrated network. For cross-timepoint validation, we simply reconstructed regulons based on one timepoint (training set) and validated the regulons on the other timepoint (test set).

For both within-timepoint and cross-timepoint cross-validations, we selected 340 sgRNAs that 1) had at least medium-efficacy (>50% knockdown) and 2) were covered by more than 15 cells at both timepoint. Out of the 250 TRs (thus 250 regulons) targeted by the 340 sgRNAs, 247 were successfully reconstructed by T1 dataset and 241 were reconstructed by T2 dataset. The unsuccessful regulon reconstruction was mainly due to low level of TR expression in single cells.

### DNase-Seq validation

DNase-Seq data were accessed from Cistrome database (Liu, Ortiz et al. 2011) on Apr 29, 2020. Context-specific data were extracted by limiting the “Cell_line” to contain “HT29”. A peak was considered to be associated with a specific gene if it overlaps with the genomic region of that gene. Peaks that did not overlap with any gene were removed.

### Protein activity inference of CCLE HT-29 and TR-perturbed HT-29

The gene expression profile of HT-29 was accessed from the Cancer Cell Line Encyclopedia (CCLE) database on Jul 10, 2020. The gene expression signature of HT-29 was then generated by z-score transformation implemented by viperSignature function from the VIPER R package, comparing the normalized expression level of HT-29 against all the other CCLE cell lines. The gene expression signatures of TR-perturbations were calculated as z-score as described in “**Differential expression analysis”.**

For all the gene expression signatures described above, the R package VIPER (Alvarez, Shen et al. 2016) was used to infer protein activity. Specifically, metaVIPER (Ding, Douglass et al. 2018) (as implement in the VIPER R package) was used to combine the information of T1 network and T2 network.

### Characterization of activity regulating network (ARN) edges (Figure 3A)

The activity regulating network was first constructed by extracting TR-TR edges with statistically significant (FDR < 0.1) differential protein activity after perturbing one of the TRs. To consolidate the network constructed at T1 and T2, we took the union of the two networks. For edges with different modes of regulation (activation or repression) at T1 and T2, we kept the T1 edge and discarded the T2 edge.

An ARN edge was characterized by “transcriptional” if the differential protein activity had a corresponding differential gene expression (FDR < 0.2) of the same TR following the same perturbation. An interaction was characterized by “pathway” if the two TRs co-exist in a “pathway” gene set. ‘‘Pathway’’ gene sets were taken from a collection of the curated gene sets (c2) from MSigDB (Liberzon, Subramanian et al. 2011, Liberzon, Birger et al. 2015). Protein-protein interactions (PPI) were taken from PrePPI (Zhang, Petrey et al. 2012), BioPlex 3.0 Network for HCT116 (Huttlin, Bruckner et al. 2020), and CORUM (Ruepp, Waegele et al. 2010). Two TRs were characterized by “overlapping regulons” if the top 50 targets (ranked by likelihood) of the two regulons have statistically significant overlap (by Fisher’s exact test, FDR < 0.1).

### Multi-layer regulatory network reconstruction

The multi-layer network (MLN) was reconstructed in the following 2 steps: 1) ARN and TRN were reconstructed separately by extracting TR-TR edges with statistically significant differential protein activity (FDR < 0.1) or different expression (FDR < 0.2) after perturbing one of the TRs; for edges with different modes of regulation (activation or repression) at T1 and T2, we kept the T1 edge and discarded the T2 edge; 2) ARN edges that had corresponding TRN edges were discarded.

### Criteria for choosing TRs to include in TReK

Beginning with a list of 2,527 TRs (Table S1), we first removed TRs that were unlikely to be actively transcribed in HT-29 using a Gaussian mixture model. The remaining 1,826 genes were filtered to include candidate COAD regulators by PanCancer VIPER analysis (*n* = 30), MOMA-inferred checkpoint regulators (Paull, Aytes et al. 2021) (*n* = 69), the union of top VIPER scores by absolute value in TCGA stomach and esophageal carcinoma (STES) and COAD (*n* = 513), CHOPD TCGA PanCancer checkpoint regulators (n = 293), and gene expression (*n* = 95).

We began with a manually curated list of 2,527 Transcriptional Regulator genes (hereafter TRs) annotated by the Gene Ontology as either transcription factors or transcription cofactors (Table S1) (Alvarez, Shen et al. 2016). Candidate genes were selected by first removing TRs whose expression was not distinguishable from noise, and then by ranking remaining TRs based on their overall transcription levels and potential role as Master Regulators in COAD, based on MOMA analysis (Paull, Aytes et al. 2021) (see Methods). Each gene was targeted by the 3 best sgRNAs predicted by (Sanson, Hanna et al. 2018).

---

**Figure S1.** Overview of TReK experimental framework. CROP-seq and related methods are based on the realization that, when using CRISPR technologies, cells can replace wells as the experimental vessel as long as we know which sgRNA(s) each cell contained. To that end, a number of strategies for capturing sgRNAs in single cells have been described (Adamson et al., 2016; Datlinger et al., 2017; Dixit et al., 2016; Hill et al., 2018; Replogle et al., 2020). In CROP-Seq, the sgRNA expression cassette is duplicated during lentiviral integration. One copy of the sgRNA, though not functional, is polyadenylated and, consequently, can be captured by oligo(dT) primers. An alternative strategy is to directly capture non-polyadenylated, functional sgRNA transcripts by incorporating primer binding sites (Capture Sequence or CaptureSeq) into the sgRNA scaffold (Replogle et al., 2020). Here we combined the two strategies by replacing the original sgRNA scaffold on CROPSeq-Guide-Puro with Capture Sequence-containing sgRNA scaffold. To conduct TReK experiment, each single cell is transduced with a single sgRNA. The resulting pool of cells is subject to single-cell RNA sequencing. Transcripts containing the sgRNA sequence are expressed by both RNA polymerase II and RNA polymerase III and are captured by oligo-dT primer and 10X Chromium CaptureSeq gel beads, respectively. The resulting whole transcriptome sequencing data is demultiplexed based on cell barcode as well as the sgRNAs that were expressed in each cell.

**Figure S2.** Constitutive and inducible dCas9-expressing vectors. The constitutive dCas9 expressing vector (B) (Addgene #122205) was constructed based on the backbone of Addgene #85969 (Adamson, Norman et al. 2016) with UCOE (ubiquitous chromatin open element) (Antoniou, Harland et al. 2003) in the upstream of the promoter and blasticidin resistance as selection marker. To improve knockdown efficacy, we incorporated the bipartite repressor dCas9-KRAB-MeCP2 (Yeo, Chavez et al. 2018) into our vector. We also constructed a doxycycline-inducible dCas9-KRAB-MeCP2 vector (A) (Addgene #140690) using a TRE promoter (Gossen, Freundlieb et al. 1995). (C) Validation of the inducible vector. Colorectal cancer cell line HT-29 was lentiviral transduced with pLX-TRE-dCas9-KRAB-MeCP-P2A-BSD. After antibiotic selection with blasticidin (20 ug/ml for 5 days), 1 ug/ml doxycycline was added and cell lysate was collected after indicated time for subsequent western blot analysis of protein abundance.

**Figure S3.** Preprocessing of TReK data. (A) Overview of preprocessing pipeline. (B) Background correction for sgRNA expression. A conceptual illustration of the distribution of log(CPM+1) of a specific sgRNA across all cells is plotted as a histogram. A 2-Gaussian Mixture Model was fitted with red curve representing the low-mean Gaussian distribution and blue curve representing the high-mean Gaussian distribution. (C) Distribution of the number of detected genes in single cells with >5,000 UMI counts. (D) Relationship between total UMI count and number of sgRNA detected in single cells. All T2 cells were grouped into 6 equally-sized bins and the fraction of unique sgRNA, multiple sgRNA, and sgRNA-undetectable (no sgRNA) cells in each bin are shown. T2 dataset was shown as a representative. (E-G) Number of RNA Polymerase II-transcribed (polyA-ed), RNA Polymerase III-transcribed (non-polyA-ed), or consolidated sgRNA transcripts detected in each single cell.

**Figure S4.** Robust knockdown achieved by short time CRISPR interference. (A,B) CRISPRi-mediated target gene depletion at pseudo-bulk sample level. For each sgRNA, the relative expression level of its target gene was calculated as described in Methods. Distribution of all sgRNAs is plotted. As a reference, a null distribution calculated from a sgRNA-shuffled dataset is also shown. (C,D) CRISPRi-mediated target gene depletion at single-cell level in T2 experiment. Log(CPM+1) of target genes expressed in all cells expressing all other sgRNA (“All Other”), cells expressing scrambled sgRNAs (“NonTargeting”), and cells expressing on-target sgRNAs are shown in violin-boxplots.

**Figure S5.** Consistency between perturbed gene expression signatures. (A,B) Pearson correlations between gene expression signatures of different sgRNAs targeting the same TR. For each perturbed TR, the highest correlation is shown among all 3 pairwise correlations (from 3 sgRNAs). High-correlation perturbations (orange dots) are defined as sgRNAs having a Pearson correlation higher than the 99 percentile of Pearson correlations between sgRNAs targeting different TRs (grey dashed line). All Pearson correlations were calculated on 2,000 genes with the highest expression level variations across all perturbations. (C-E). Comparison of knockdown efficacy (C), number of differentially expressed genes (D), and number of cells (E) of high-correlation and low-correlation perturbations. Knockdown efficacy is shown as log2 fold change of target TR expression. Statistical significance was calculated using the two-tailed Mann-Whitney U test. (F) Protein half-lives of early (T1) high-correlation TRs and late (T2) high-correlation TRs. Protein half-lives were measured in hours by mass spectrometry (Cambridge, Gnad et al. 2011). Statistical significance was calculated using the one-tailed Mann-Whitney U test. (G,H) Consistency between genetically perturbed bulk sequencing and TReK perturbed transcriptomes. TReK gene expression signatures were calculated as described in Methods. In each GSEA plots, the upper gene set represents upregulated genes identified in bulk sequencing and the lower gene set represents downregulated genes. Bulk sequencing differentially expressed genes are taken from Wenzel et al., Oncogene 2020 by FDR=1x10^-5^ as threshold.

**Figure S6.** Cell number saturation analysis of network refinement. 7 TRs are shown here selected based on two criteria: 1) TRs with statistically significant (*p* < 0.05) inactive activities in the T1 5-fold cross-validation; 2) TRs perturbed in more than 60 cells in T1 5-fold cross-validation dataset. At each cell number, 20 sub-datasets were randomly down-sampled for 5-fold cross-validation and the median Stouffer’s method-integrated protein activities are shown.

**Figure S7.** Transcriptional and post-transcriptional interactions in multi-layer network. Intersections of different types of interactions in TRN and ARN interactions before (A) and after (B) removing ARN interactions with corresponding TRN interaction.

**Figure S8.** Correlation between perturbation efficacy and enrichment of downstream transcriptional targets. (A) Correlation between *CTNNB1* perturbation efficacy and enrichment of Wnt-β-catenin pathway genes. (B) Correlation between *MYBL2* perturbation efficacy and enrichment of G2M checkpoint genes.

**Figure S9.** Enrichment analysis of E2F targets after *E2F7* perturbation at different timepoints. Gene set enrichment analysis of E2F targets on gene expression signatures after 3 different *E2F7*-targeting sgRNA perturbations. Empirical p-values are shown.

**Figure S10.** Identification of upstream regulators and co-factors of MYC. (A) Log(CPM+1) of *MAX* expressed in all cells expressing sgRNA targeting other genes (“All Other”), cells expressing scrambled sgRNAs (“NonTargeting”), and cells expressing *MAX*-targeting sgRNAs shown in violin-boxplots. (B-D) GSEA of MYC targets after different *MAX*-targeting sgRNA perturbations. Empirical p-values are shown. (E) Venn diagram showing the overlap between TReK-predicted MYC modulators and CINDY-predicted MYC modulators. Statistical significance of the overlap was calculated using the Fisher’s exact test. (F) Venn diagram showing MYC modulators, identified by both TReK and CINDY, with different mechanistic charaterizations.

## Notes

### Competing Interest Statement

A.C. is founder, equity holder, consultant, and director of DarwinHealth Inc., a company that has licensed some of the algorithms used in this manuscript from Columbia University. DarwinHealth Inc. has licensed IP related to these algorithms from Columbia University. Columbia University is an equity holder in DarwinHealth Inc.

## References

Adamson, B., T. M. Norman, M. Jost, M. Y. Cho, J. K. Nunez, Y. Chen, J. E. Villalta, L. A. Gilbert, M. A. Horlbeck, M. Y. Hein, R. A. Pak, A. N. Gray, C. A. Gross, A. Dixit, O. Parnas, A. Regev and J. S. Weissman (2016). “A Multiplexed Single-Cell CRISPR Screening Platform Enables Systematic Dissection of the Unfolded Protein Response.” Cell 167(7): 1867–1882 e1821.

Aibar, S., C. B. Gonzalez-Blas, T. Moerman, V. A. Huynh-Thu, H. Imrichova, G. Hulselmans, F. Rambow, J. C. Marine, P. Geurts, J. Aerts, J. van den Oord, Z. K. Atak, J. Wouters and S. Aerts (2017). “SCENIC: single-cell regulatory network inference and clustering.” Nat Methods 14(11): 1083–1086.

Alon, U. (2007). “Network motifs: theory and experimental approaches.” Nat Rev Genet 8(6): 450–461.

Alvarez, M. J., Y. Shen, F. M. Giorgi, A. Lachmann, B. B. Ding, B. H. Ye and A. Califano (2016). “Functional characterization of somatic mutations in cancer using network-based inference of protein activity.” Nat Genet 48(8): 838–847.

Antoniou, M., L. Harland, T. Mustoe, S. Williams, J. Holdstock, E. Yague, T. Mulcahy, M. Griffiths, S. Edwards, P. A. Ioannou, A. Mountain and R. Crombie (2003). “Transgenes encompassing dual-promoter CpG islands from the human TBP and HNRPA2B1 loci are resistant to heterochromatin-mediated silencing.” Genomics 82(3): 269–279.

Arumugam, K., W. Shin, V. Schiavone, L. Vlahos, X. Tu, D. Carnevali, J. Kesner, E. O. Paull, N. Romo, P. Subramaniam, J. Worley, X. Tan, A. Califano and M. P. Cosma (2020). “The Master Regulator Protein BAZ2B Can Reprogram Human Hematopoietic Lineage-Committed Progenitors into a Multipotent State.” Cell Rep 33(10): 108474.

Aytes, A., A. Mitrofanova, C. Lefebvre, M. J. Alvarez, M. Castillo-Martin, T. Zheng, J. A. Eastham, A. Gopalan, K. J. Pienta, M. M. Shen, A. Califano and C. Abate-Shen (2014). “Cross-species regulatory network analysis identifies a synergistic interaction between FOXM1 and CENPF that drives prostate cancer malignancy.” Cancer Cell 25(5): 638–651.

Basso, K., A. A. Margolin, G. Stolovitzky, U. Klein, R. Dalla-Favera and A. Califano (2005). “Reverse engineering of regulatory networks in human B cells.” Nat Genet 37(4): 382–390.

Boorsma, A., X. J. Lu, A. Zakrzewska, F. M. Klis and H. J. Bussemaker (2008). “Inferring condition-specific modulation of transcription factor activity in yeast through regulon-based analysis of genomewide expression.” PLoS One 3(9): e3112.

Brantjes, H., N. Barker, J. van Es and H. Clevers (2002). “TCF: Lady Justice casting the final verdict on the outcome of Wnt signalling.” Biol Chem 383(2): 255–261.

Cambridge, S. B., F. Gnad, C. Nguyen, J. L. Bermejo, M. Kruger and M. Mann (2011). “Systems-wide proteomic analysis in mammalian cells reveals conserved, functional protein turnover.” J Proteome Res 10(12): 5275–5284.

Cancer Genome Atlas, N. (2012). “Comprehensive molecular characterization of human colon and rectal cancer.” Nature 487(7407): 330–337.

Carro, M. S., W. K. Lim, M. J. Alvarez, R. J. Bollo, X. Zhao, E. Y. Snyder, E. P. Sulman, S. L. Anne, F. Doetsch, H. Colman, A. Lasorella, K. Aldape, A. Califano and A. Iavarone (2010). “The transcriptional network for mesenchymal transformation of brain tumours.” Nature 463(7279): 318–325.

Chen, J. C., M. J. Alvarez, F. Talos, H. Dhruv, G. E. Rieckhof, A. Iyer, K. L. Diefes, K. Aldape, M. Berens, M. M. Shen and A. Califano (2014). “Identification of causal genetic drivers of human disease through systems-level analysis of regulatory networks.” Cell 159(2): 402–414.

Cragg, J. G. (1971). “Some Statistical Models for Limited Dependent Variables with Application to the Demand for Durable Goods.” Econometrica 39(5): 16.

Dang, C. V. (2012). “MYC on the path to cancer.” Cell 149(1): 22–35.

Datlinger, P., A. F. Rendeiro, T. Boenke, M. Senekowitsch, T. Krausgruber, D. Barreca and C. Bock (2021). “Ultra-high-throughput single-cell RNA sequencing and perturbation screening with combinatorial fluidic indexing.” Nat Methods 18(6): 635–642.

Datlinger, P., A. F. Rendeiro, C. Schmidl, T. Krausgruber, P. Traxler, J. Klughammer, L. C. Schuster, A. Kuchler, D. Alpar and C. Bock (2017). “Pooled CRISPR screening with single-cell transcriptome readout.” Nat Methods 14(3): 297–301.

Davidson, E. H., J. P. Rast, P. Oliveri, A. Ransick, C. Calestani, C. H. Yuh, T. Minokawa, G. Amore, V. Hinman, C. Arenas-Mena, O. Otim, C. T. Brown, C. B. Livi, P. Y. Lee, R. Revilla, A. G. Rust, Z. Pan, M. J. Schilstra, P. J. Clarke, M. I. Arnone, L. Rowen, R. A. Cameron, D. R. McClay, L. Hood and H. Bolouri (2002). “A genomic regulatory network for development.” Science 295(5560): 1669–1678.

de Bruin, A., B. Maiti, L. Jakoi, C. Timmers, R. Buerki and G. Leone (2003). “Identification and characterization of E2F7, a novel mammalian E2F family member capable of blocking cellular proliferation.” J Biol Chem 278(43): 42041–42049.

Di Stefano, L., M. R. Jensen and K. Helin (2003). “E2F7, a novel E2F featuring DP-independent repression of a subset of E2F-regulated genes.” EMBO J 22(23): 6289–6298.

Ding, H., E. F. Douglass, Jr., A. M. Sonabend, A. Mela, S. Bose, C. Gonzalez, P. D. Canoll, P. A. Sims, M. J. Alvarez and A. Califano (2018). “Quantitative assessment of protein activity in orphan tissues and single cells using the metaVIPER algorithm.” Nat Commun 9(1): 1471.

Dixit, A., O. Parnas, B. Li, J. Chen, C. P. Fulco, L. Jerby-Arnon, N. D. Marjanovic, D. Dionne, T. Burks, R. Raychowdhury, B. Adamson, T. M. Norman, E. S. Lander, J. S. Weissman, N. Friedman and A. Regev (2016). “Perturb-Seq: Dissecting Molecular Circuits with Scalable Single-Cell RNA Profiling of Pooled Genetic Screens.” Cell 167(7): 1853–1866 e1817.

Dutta, A., C. Le Magnen, A. Mitrofanova, X. Ouyang, A. Califano and C. Abate-Shen (2016). “Identification of an NKX3.1-G9a-UTY transcriptional regulatory network that controls prostate differentiation.” Science 352(6293): 1576–1580.

Finak, G., A. McDavid, M. Yajima, J. Deng, V. Gersuk, A. K. Shalek, C. K. Slichter, H. W. Miller, M. J. McElrath, M. Prlic, P. S. Linsley and R. Gottardo (2015). “MAST: a flexible statistical framework for assessing transcriptional changes and characterizing heterogeneity in single-cell RNA sequencing data.” Genome Biol 16: 278.

Ghandi, M., F. W. Huang, J. Jane-Valbuena, G. V. Kryukov, C. C. Lo, E. R. McDonald, 3rd, J. Barretina, E. T. Gelfand, C. M. Bielski, H. Li, K. Hu, A. Y. Andreev-Drakhlin, J. Kim, J. M. Hess, B. J. Haas, F. Aguet, B. A. Weir, M. V. Rothberg, B. R. Paolella, M. S. Lawrence, R. Akbani, Y. Lu, H. L. Tiv, P. C. Gokhale, A. de Weck, A. A. Mansour, C. Oh, J. Shih, K. Hadi, Y. Rosen, J. Bistline, K. Venkatesan, A. Reddy, D. Sonkin, M. Liu, J. Lehar, J. M. Korn, D. A. Porter, M. D. Jones, J. Golji, G. Caponigro, J. E. Taylor, C. M. Dunning, A. L. Creech, A. C. Warren, J. M. McFarland, M. Zamanighomi, A. Kauffmann, N. Stransky, M. Imielinski, Y. E. Maruvka, A. D. Cherniack, A. Tsherniak, F. Vazquez, J. D. Jaffe, A. A. Lane, D. M. Weinstock, C. M. Johannessen, M. P. Morrissey, F. Stegmeier, R. Schlegel, W. C. Hahn, G. Getz, G. B. Mills, J. S. Boehm, T. R. Golub, L. A. Garraway and W. R. Sellers (2019). “Next-generation characterization of the Cancer Cell Line Encyclopedia.” Nature 569(7757): 503–508.

Gilbert, L. A., M. A. Horlbeck, B. Adamson, J. E. Villalta, Y. Chen, E. H. Whitehead, C. Guimaraes, B. Panning, H. L. Ploegh, M. C. Bassik, L. S. Qi, M. Kampmann and J. S. Weissman (2014). “Genome-Scale CRISPR-Mediated Control of Gene Repression and Activation.” Cell 159(3): 647–661.

Giorgi, F. M., G. Lopez, J. H. Woo, B. Bisikirska, A. Califano and M. Bansal (2014). “Inferring protein modulation from gene expression data using conditional mutual information.” PLoS One 9(10): e109569.

Gossen, M., S. Freundlieb, G. Bender, G. Muller, W. Hillen and H. Bujard (1995). “Transcriptional activation by tetracyclines in mammalian cells.” Science 268(5218): 1766–1769.

Hill, A. J., J. L. McFaline-Figueroa, L. M. Starita, M. J. Gasperini, K. A. Matreyek, J. Packer, D. Jackson, J. Shendure and C. Trapnell (2018). “On the design of CRISPR-based single-cell molecular screens.” Nat Methods 15(4): 271–274.

Hironaka, K., V. M. Factor, D. F. Calvisi, E. A. Conner and S. S. Thorgeirsson (2003). “Dysregulation of DNA repair pathways in a transforming growth factor alpha/c-myc transgenic mouse model of accelerated hepatocarcinogenesis.” Lab Invest 83(5): 643–654.

Huttlin, E. L., R. J. Bruckner, J. Navarrete-Perea, J. R. Cannon, K. Baltier, F. Gebreab, M. P. Gygi, A. Thornock, G. Zarraga, S. Tam, J. Szpyt, A. Panov, H. Parzen, S. Fu, A. Golbazi, E. Maenpaa, K. Stricker, S. G. Thakurta, R. Rad, J. Pan, D. P. Nusinow, J. A. Paulo, D. K. Schweppe, L. P. Vaites, J. W. Harper and S. P. Gygi (2020). “Dual Proteome-scale Networks Reveal Cell-specific Remodeling of the Human Interactome.” 2020.2001.2019.905109.

Huynh-Thu, V. A., A. Irrthum, L. Wehenkel and P. Geurts (2010). “Inferring regulatory networks from expression data using tree-based methods.” PLoS One 5(9).

Hwang, S., C. Y. Kim, S. Yang, E. Kim, T. Hart, E. M. Marcotte and I. Lee (2019). “HumanNet v2: human gene networks for disease research.” Nucleic Acids Res 47(D1): D573–D580.

Karlsson, A., D. Deb-Basu, A. Cherry, S. Turner, J. Ford and D. W. Felsher (2003). “Defective double-strand DNA break repair and chromosomal translocations by MYC overexpression.” Proc Natl Acad Sci U S A 100(17): 9974–9979.

Khurana, E., Y. Fu, J. Chen and M. Gerstein (2013). “Interpretation of genomic variants using a unified biological network approach.” PLoS Comput Biol 9(3): e1002886.

Liberzon, A., C. Birger, H. Thorvaldsdottir, M. Ghandi, J. P. Mesirov and P. Tamayo (2015). “The Molecular Signatures Database (MSigDB) hallmark gene set collection.” Cell Syst 1(6): 417–425.

Liberzon, A., A. Subramanian, R. Pinchback, H. Thorvaldsdottir, P. Tamayo and J. P. Mesirov (2011). “Molecular signatures database (MSigDB) 3.0.” Bioinformatics 27(12): 1739–1740.

Liu, T., J. A. Ortiz, L. Taing, C. A. Meyer, B. Lee, Y. Zhang, H. Shin, S. S. Wong, J. Ma, Y. Lei, U. J. Pape, M. Poidinger, Y. Chen, K. Yeung, M. Brown, Y. Turpaz and X. S. Liu (2011). “Cistrome: an integrative platform for transcriptional regulation studies.” Genome Biol 12(8): R83.

Ma, L., J. Wagner, J. J. Rice, W. Hu, A. J. Levine and G. A. Stolovitzky (2005). “A plausible model for the digital response of p53 to DNA damage.” Proc Natl Acad Sci U S A 102(40): 14266–14271.

Margolin, A. A., I. Nemenman, K. Basso, C. Wiggins, G. Stolovitzky, R. Dalla Favera and A. Califano (2006). “ARACNE: an algorithm for the reconstruction of gene regulatory networks in a mammalian cellular context.” BMC Bioinformatics 7 **Suppl 1**: S7.

Mei, S., Q. Qin, Q. Wu, H. Sun, R. Zheng, C. Zang, M. Zhu, J. Wu, X. Shi, L. Taing, T. Liu, M. Brown, C. A. Meyer and X. S. Liu (2017). “Cistrome Data Browser: a data portal for ChIP-Seq and chromatin accessibility data in human and mouse.” Nucleic Acids Res 45(D1): D658–D662.

Murray, J. I. (2018). “Systems biology of embryonic development: Prospects for a complete understanding of the Caenorhabditis elegans embryo.” Wiley Interdiscip Rev Dev Biol 7(3): e314.

Obradovic, A., N. Chowdhury, S. M. Haake, C. Ager, V. Wang, L. Vlahos, X. V. Guo, D. H. Aggen, W. K. Rathmell, E. Jonasch, J. E. Johnson, M. Roth, K. E. Beckermann, B. I. Rini, J. McKiernan, A. Califano and C. G. Drake (2021). “Single-cell protein activity analysis identifies recurrence-associated renal tumor macrophages.” Cell 184(11): 2988–3005 e2916.

Paull, E. O., A. Aytes, S. J. Jones, P. S. Subramaniam, F. M. Giorgi, E. F. Douglass, S. Tagore, B. Chu, A. Vasciaveo, S. Zheng, R. Verhaak, C. Abate-Shen, M. J. Alvarez and A. Califano (2021). “A modular master regulator landscape controls cancer transcriptional identity.” Cell 184(2): 334–351 e320.

Pe’er, D. (2005). “Bayesian network analysis of signaling networks: a primer.” Sci STKE 2005(281): pl4.

Rajbhandari, P., G. Lopez, C. Capdevila, B. Salvatori, J. Yu, R. Rodriguez-Barrueco, D. Martinez, M. Yarmarkovich, N. Weichert-Leahey, B. J. Abraham, M. J. Alvarez, A. Iyer, J. L. Harenza, D. Oldridge, K. De Preter, J. Koster, S. Asgharzadeh, R. C. Seeger, J. S. Wei, J. Khan, J. Vandesompele, P. Mestdagh, R. Versteeg, A. T. Look, R. A. Young, A. Iavarone, A. Lasorella, J. M. Silva, J. M. Maris and A. Califano (2018). “Cross-Cohort Analysis Identifies a TEAD4-MYCN Positive Feedback Loop as the Core Regulatory Element of High-Risk Neuroblastoma.” Cancer Discov 8(5): 582–599.

Rennoll, S. and G. Yochum (2015). “Regulation of MYC gene expression by aberrant Wnt/beta-catenin signaling in colorectal cancer.” World J Biol Chem 6(4): 290–300.

Replogle, J. M., T. M. Norman, A. Xu, J. A. Hussmann, J. Chen, J. Z. Cogan, E. J. Meer, J. M. Terry, D. P. Riordan, N. Srinivas, I. T. Fiddes, J. G. Arthur, L. J. Alvarado, K. A. Pfeiffer, T. S. Mikkelsen, J. S. Weissman and B. Adamson (2020). “Combinatorial single-cell CRISPR screens by direct guide RNA capture and targeted sequencing.” Nat Biotechnol 38(8): 954–961.

Rolland, T., M. Tasan, B. Charloteaux, S. J. Pevzner, Q. Zhong, N. Sahni, S. Yi, I. Lemmens, C. Fontanillo, R. Mosca, A. Kamburov, S. D. Ghiassian, X. Yang, L. Ghamsari, D. Balcha, B. E. Begg, P. Braun, M. Brehme, M. P. Broly, A. R. Carvunis, D. Convery-Zupan, R. Corominas, J. Coulombe-Huntington, E. Dann, M. Dreze, A. Dricot, C. Fan, E. Franzosa, F. Gebreab, B. J. Gutierrez, M. F. Hardy, M. Jin, S. Kang, R. Kiros, G. N. Lin, K. Luck, A. MacWilliams, J. Menche, R. R. Murray, A. Palagi, M. M. Poulin, X. Rambout, J. Rasla, P. Reichert, V. Romero, E. Ruyssinck, J. M. Sahalie, A. Scholz, A. A. Shah, A. Sharma, Y. Shen, K. Spirohn, S. Tam, A. O. Tejeda, S. A. Trigg, J. C. Twizere, K. Vega, J. Walsh, M. E. Cusick, Y. Xia, A. L. Barabasi, L. M. Iakoucheva, P. Aloy, J. De Las Rivas, J. Tavernier, M. A. Calderwood, D. E. Hill, T. Hao, F. P. Roth and M. Vidal (2014). “A proteome-scale map of the human interactome network.” Cell 159(5): 1212–1226.

Rual, J. F., K. Venkatesan, T. Hao, T. Hirozane-Kishikawa, A. Dricot, N. Li, G. F. Berriz, F. D. Gibbons, M. Dreze, N. Ayivi-Guedehoussou, N. Klitgord, C. Simon, M. Boxem, S. Milstein, J. Rosenberg, D. S. Goldberg, L. V. Zhang, S. L. Wong, G. Franklin, S. Li, J. S. Albala, J. Lim, C. Fraughton, E. Llamosas, S. Cevik, C. Bex, P. Lamesch, R. S. Sikorski, J. Vandenhaute, H. Y. Zoghbi, A. Smolyar, S. Bosak, R. Sequerra, L. Doucette-Stamm, M. E. Cusick, D. E. Hill, F. P. Roth and M. Vidal (2005). “Towards a proteome-scale map of the human protein-protein interaction network.” Nature 437(7062): 1173–1178.

Ruepp, A., B. Waegele, M. Lechner, B. Brauner, I. Dunger-Kaltenbach, G. Fobo, G. Frishman, C. Montrone and H. W. Mewes (2010). “CORUM: the comprehensive resource of mammalian protein complexes--2009.” Nucleic Acids Res 38(Database issue): D497-501.

Sablitzky, F., A. Moore, M. Bromley, R. W. Deed, J. S. Newton and J. D. Norton (1998). “Stage- and subcellular-specific expression of Id proteins in male germ and Sertoli cells implicates distinctive regulatory roles for Id proteins during meiosis, spermatogenesis, and Sertoli cell function.” Cell Growth Differ 9(12): 1015–1024.

Sanson, K. R., R. E. Hanna, M. Hegde, K. F. Donovan, C. Strand, M. E. Sullender, E. W. Vaimberg, A. Goodale, D. E. Root, F. Piccioni and J. G. Doench (2018). “Optimized libraries for CRISPR-Cas9 genetic screens with multiple modalities.” Nat Commun 9(1): 5416.

Segeren, H. A., L. M. van Rijnberk, E. Moreno, F. M. Riemers, E. A. van Liere, R. Yuan, R. Wubbolts, A. de Bruin and B. Westendorp (2020). “Excessive E2F Transcription in Single Cancer Cells Precludes Transient Cell-Cycle Exit after DNA Damage.” Cell Rep 33(9): 108449.

Smith, V. A., J. Yu, T. V. Smulders, A. J. Hartemink and E. D. Jarvis (2006). “Computational inference of neural information flow networks.” PLoS Comput Biol 2(11): e161.

Soneson, C. and M. D. Robinson (2018). “Bias, robustness and scalability in single-cell differential expression analysis.” Nat Methods 15(4): 255–261.

Subramanian, A., P. Tamayo, V. K. Mootha, S. Mukherjee, B. L. Ebert, M. A. Gillette, A. Paulovich, S. L. Pomeroy, T. R. Golub, E. S. Lander and J. P. Mesirov (2005). “Gene set enrichment analysis: a knowledge-based approach for interpreting genome-wide expression profiles.” Proc Natl Acad Sci U S A 102(43): 15545–15550.

Takahashi, J. S. (2017). “Transcriptional architecture of the mammalian circadian clock.” Nat Rev Genet 18(3): 164–179.

Tate, J. G., S. Bamford, H. C. Jubb, Z. Sondka, D. M. Beare, N. Bindal, H. Boutselakis, C. G. Cole, C. Creatore, E. Dawson, P. Fish, B. Harsha, C. Hathaway, S. C. Jupe, C. Y. Kok, K. Noble, L. Ponting, C. C. Ramshaw, C. E. Rye, H. E. Speedy, R. Stefancsik, S. L. Thompson, S. Wang, S. Ward, P. J. Campbell and S. A. Forbes (2019). “COSMIC: the Catalogue Of Somatic Mutations In Cancer.” Nucleic Acids Res 47(D1): D941–D947.

Vu, T. and P. K. Datta (2017). “Regulation of EMT in Colorectal Cancer: A Culprit in Metastasis.” Cancers (Basel) 9(12).

Wang, K., M. Saito, B. C. Bisikirska, M. J. Alvarez, W. K. Lim, P. Rajbhandari, Q. Shen, I. Nemenman, K. Basso, A. A. Margolin, U. Klein, R. Dalla-Favera and A. Califano (2009). “Genome-wide identification of post-translational modulators of transcription factor activity in human B cells.” Nat Biotechnol 27(9): 829–839.

Wenzel, J., K. Rose, E. B. Haghighi, C. Lamprecht, G. Rauen, V. Freihen, R. Kesselring, M. Boerries and A. Hecht (2020). “Loss of the nuclear Wnt pathway effector TCF7L2 promotes migration and invasion of human colorectal cancer cells.” Oncogene 39(19): 3893–3909.

Westendorp, B., M. Mokry, M. J. Groot Koerkamp, F. C. Holstege, E. Cuppen and A. de Bruin (2012). “E2F7 represses a network of oscillating cell cycle genes to control S-phase progression.” Nucleic Acids Res 40(8): 3511–3523.

Woo, J. H., Y. Shimoni, W. S. Yang, P. Subramaniam, A. Iyer, P. Nicoletti, M. Rodriguez Martinez, G. Lopez, M. Mattioli, R. Realubit, C. Karan, B. R. Stockwell, M. Bansal and A. Califano (2015). “Elucidating Compound Mechanism of Action by Network Perturbation Analysis.” Cell 162(2): 441–451.

Yagi, K., M. Furuhashi, H. Aoki, D. Goto, H. Kuwano, K. Sugamura, K. Miyazono and M. Kato (2002). “c-myc is a downstream target of the Smad pathway.” J Biol Chem 277(1): 854–861.

Yeo, N. C., A. Chavez, A. Lance-Byrne, Y. Chan, D. Menn, D. Milanova, C. C. Kuo, X. Guo, S. Sharma, A. Tung, R. J. Cecchi, M. Tuttle, S. Pradhan, E. T. Lim, N. Davidsohn, M. R. Ebrahimkhani, J. J. Collins, N. E. Lewis, S. Kiani and G. M. Church (2018). “An enhanced CRISPR repressor for targeted mammalian gene regulation.” Nat Methods 15(8): 611–616.

Yue, M., J. Jiang, P. Gao, H. Liu and G. Qing (2017). “Oncogenic MYC Activates a Feedforward Regulatory Loop Promoting Essential Amino Acid Metabolism and Tumorigenesis.” Cell Rep 21(13): 3819–3832.

Zhang, Q. C., D. Petrey, L. Deng, L. Qiang, Y. Shi, C. A. Thu, B. Bisikirska, C. Lefebvre, D. Accili, T. Hunter, T. Maniatis, A. Califano and B. Honig (2012). “Structure-based prediction of protein-protein interactions on a genome-wide scale.” Nature 490(7421): 556–560.

Zhang, Q. C., D. Petrey, J. I. Garzon, L. Deng and B. Honig (2013). “PrePPI: a structure-informed database of protein-protein interactions.” Nucleic Acids Res 41(Database issue): D828-833.

Zheng, R., C. Wan, S. Mei, Q. Qin, Q. Wu, H. Sun, C. H. Chen, M. Brown, X. Zhang, C. A. Meyer and X. S. Liu (2019). “Cistrome Data Browser: expanded datasets and new tools for gene regulatory analysis.” Nucleic Acids Res 47(D1): D729–D735.

