## Supplementary material for "Interrogation of genome-wide, experimentally dissected gene regulatory networks reveals mechanisms underlying dynamic cellular state control": Table_S5_KnownRegulationsIdentifiedByTREK.docx

**Supplementary Information**

**Table S5. Known Wnt/TGF pathway regulations identified by TREK**

| **Regulator** | **Cancer hallmark pathway** | **Evidence** |
| --- | --- | --- |
| MYC | E2F_TARGETS | (Leung, Ehmann et al. 2008) |
| MYC | G2M_CHECKPOINT | (Felsher, Zetterberg et al. 2000, Sheen, Woo et al. 2003, Yang, Xue et al. 2018) |
| MYC | MTORC1_SIGNALING | (Yue, Jiang et al. 2017) |
| MYC | MYC_TARGETS_V2 | (Liberzon, Subramanian et al. 2011, Liberzon, Birger et al. 2015) |
| MYC | DNA_REPAIR | (Hironaka, Factor et al. 2003, Karlsson, Deb-Basu et al. 2003) |
| ID2 | SPERMATOGENESIS | (Sablitzky, Moore et al. 1998) |
| ID1 | E2F_TARGETS | (Li, Xu et al. 2016) |
| HDAC2 | WNT_BETA_CATENIN_SIGNALING | (Ye, Chen et al. 2009, Gotze, Coersmeyer et al. 2014) |
| HDAC2 | E2F_TARGETS | (Brehm, Miska et al. 1998, Noh, Jung et al. 2011) |
| ID1 | MYC_TARGETS_V1 | (Qian, Lee et al. 2010, Sharma, Kolhe et al. 2016) |
| TCF7 | MYC_TARGETS_V1 | (Roose, Huls et al. 1999, Liu, Sun et al. 2019) |
| TCF7 | WNT_BETA_CATENIN_SIGNALING | (Roose, Huls et al. 1999, Kolligs, Bommer et al. 2002) |
| CTNNB1 | MYC_TARGETS_V1 | (Rennoll and Yochum 2015) |
| CTNNB1 | TNFA_SIGNALING_VIA_NFKB | (Deng, Miller et al. 2002) |
| CTNNB1 | MITOTIC_SPINDLE | (Kaplan, Meigs et al. 2004) |
| CTNNB1 | WNT_BETA_CATENIN_SIGNALING | (Liberzon, Subramanian et al. 2011, Liberzon, Birger et al. 2015) |
| CTNNB1 | MYOGENESIS | (Cui, Li et al. 2019) |
| CTNNB1 | PI3K_AKT_MTOR_SIGNALING | (Wang, Zhou et al. 2018) |
| CTNNB1 | EPITHELIAL_MESENCHYMAL_TRANSITION | (Kim, Kwon et al. 2019) |
| JUNB | MYC_TARGETS_V1 | (Vartanian, Masri et al. 2011) |
| SMAD3 | HEDGEHOG_SIGNALING | (Dennler, Andre et al. 2007) |
| TP53 | OXIDATIVE_PHOPHORYLATION | (Matoba, Kang et al. 2006, Zhang, Lin et al. 2011) |
| TP53 | REACTIVE_OXYGEN_SPECIES_PATHWAY | (Johnson, Yu et al. 1996) |
| CDK9 | APOPTOSIS | (Rahaman, Lam et al. 2019) |

**References**

Brehm, A., E. A. Miska, D. J. McCance, J. L. Reid, A. J. Bannister and T. Kouzarides (1998). "Retinoblastoma protein recruits histone deacetylase to repress transcription." Nature **391**(6667): 597-601.

Cui, S., L. Li, R. T. Yu, M. Downes, R. M. Evans, J. A. Hulin, H. P. Makarenkova and R. Meech (2019). "beta-Catenin is essential for differentiation of primary myoblasts via cooperation with MyoD and alpha-catenin." Development **146**(6).

Deng, J., S. A. Miller, H. Y. Wang, W. Xia, Y. Wen, B. P. Zhou, Y. Li, S. Y. Lin and M. C. Hung (2002). "beta-catenin interacts with and inhibits NF-kappa B in human colon and breast cancer." Cancer Cell **2**(4): 323-334.

Dennler, S., J. Andre, I. Alexaki, A. Li, T. Magnaldo, P. ten Dijke, X. J. Wang, F. Verrecchia and A. Mauviel (2007). "Induction of sonic hedgehog mediators by transforming growth factor-beta: Smad3-dependent activation of Gli2 and Gli1 expression in vitro and in vivo." Cancer Res **67**(14): 6981-6986.

Felsher, D. W., A. Zetterberg, J. Zhu, T. Tlsty and J. M. Bishop (2000). "Overexpression of MYC causes p53-dependent G2 arrest of normal fibroblasts." Proc Natl Acad Sci U S A **97**(19): 10544-10548.

Gotze, S., M. Coersmeyer, O. Muller and S. Sievers (2014). "Histone deacetylase inhibitors induce attenuation of Wnt signaling and TCF7L2 depletion in colorectal carcinoma cells." Int J Oncol **45**(4): 1715-1723.

Johnson, T. M., Z. X. Yu, V. J. Ferrans, R. A. Lowenstein and T. Finkel (1996). "Reactive oxygen species are downstream mediators of p53-dependent apoptosis." Proc Natl Acad Sci U S A **93**(21): 11848-11852.

Kaplan, D. D., T. E. Meigs, P. Kelly and P. J. Casey (2004). "Identification of a role for beta-catenin in the establishment of a bipolar mitotic spindle." J Biol Chem **279**(12): 10829-10832.

Kim, W. K., Y. Kwon, M. Jang, M. Park, J. Kim, S. Cho, D. G. Jang, W. B. Lee, S. H. Jung, H. J. Choi, B. S. Min, T. Il Kim, S. P. Hong, Y. K. Paik and H. Kim (2019). "beta-catenin activation down-regulates cell-cell junction-related genes and induces epithelial-to-mesenchymal transition in colorectal cancers." Sci Rep **9**(1): 18440.

Kolligs, F. T., G. Bommer and B. Goke (2002). "Wnt/beta-catenin/tcf signaling: a critical pathway in gastrointestinal tumorigenesis." Digestion **66**(3): 131-144.

Leung, J. Y., G. L. Ehmann, P. H. Giangrande and J. R. Nevins (2008). "A role for Myc in facilitating transcription activation by E2F1." Oncogene **27**(30): 4172-4179.

Li, B., W. W. Xu, X. Y. Guan, Y. R. Qin, S. Law, N. P. Lee, K. T. Chan, P. Y. Tam, Y. Y. Li, K. W. Chan, H. F. Yuen, S. W. Tsao, Q. Y. He and A. L. Cheung (2016). "Competitive Binding Between Id1 and E2F1 to Cdc20 Regulates E2F1 Degradation and Thymidylate Synthase Expression to Promote Esophageal Cancer Chemoresistance." Clin Cancer Res **22**(5): 1243-1255.

Liberzon, A., C. Birger, H. Thorvaldsdottir, M. Ghandi, J. P. Mesirov and P. Tamayo (2015). "The Molecular Signatures Database (MSigDB) hallmark gene set collection." Cell Syst **1**(6): 417-425.

Liberzon, A., A. Subramanian, R. Pinchback, H. Thorvaldsdottir, P. Tamayo and J. P. Mesirov (2011). "Molecular signatures database (MSigDB) 3.0." Bioinformatics **27**(12): 1739-1740.

Liu, Z., R. Sun, X. Zhang, B. Qiu, T. Chen, Z. Li, Y. Xu and Z. Zhang (2019). "Transcription factor 7 promotes the progression of perihilar cholangiocarcinoma by inducing the transcription of c-Myc and FOS-like antigen 1." EBioMedicine **45**: 181-191.

Matoba, S., J. G. Kang, W. D. Patino, A. Wragg, M. Boehm, O. Gavrilova, P. J. Hurley, F. Bunz and P. M. Hwang (2006). "p53 regulates mitochondrial respiration." Science **312**(5780): 1650-1653.

Noh, J. H., K. H. Jung, J. K. Kim, J. W. Eun, H. J. Bae, H. J. Xie, Y. G. Chang, M. G. Kim, W. S. Park, J. Y. Lee and S. W. Nam (2011). "Aberrant regulation of HDAC2 mediates proliferation of hepatocellular carcinoma cells by deregulating expression of G1/S cell cycle proteins." PLoS One **6**(11): e28103.

Qian, T., J. Y. Lee, J. H. Park, H. J. Kim and G. Kong (2010). "Id1 enhances RING1b E3 ubiquitin ligase activity through the Mel-18/Bmi-1 polycomb group complex." Oncogene **29**(43): 5818-5827.

Rahaman, M. H., F. Lam, L. Zhong, T. Teo, J. Adams, M. Yu, R. W. Milne, C. Pepper, N. A. Lokman, C. Ricciardelli, M. K. Oehler and S. Wang (2019). "Targeting CDK9 for treatment of colorectal cancer." Mol Oncol **13**(10): 2178-2193.

Rennoll, S. and G. Yochum (2015). "Regulation of MYC gene expression by aberrant Wnt/beta-catenin signaling in colorectal cancer." World J Biol Chem **6**(4): 290-300.

Roose, J., G. Huls, M. van Beest, P. Moerer, K. van der Horn, R. Goldschmeding, T. Logtenberg and H. Clevers (1999). "Synergy between tumor suppressor APC and the beta-catenin-Tcf4 target Tcf1." Science **285**(5435): 1923-1926.

Sablitzky, F., A. Moore, M. Bromley, R. W. Deed, J. S. Newton and J. D. Norton (1998). "Stage- and subcellular-specific expression of Id proteins in male germ and Sertoli cells implicates distinctive regulatory roles for Id proteins during meiosis, spermatogenesis, and Sertoli cell function." Cell Growth Differ **9**(12): 1015-1024.

Sharma, B. K., R. Kolhe, S. M. Black, J. R. Keller, N. F. Mivechi and A. Satyanarayana (2016). "Inhibitor of differentiation 1 transcription factor promotes metabolic reprogramming in hepatocellular carcinoma cells." FASEB J **30**(1): 262-275.

Sheen, J. H., J. K. Woo and R. B. Dickson (2003). "c-Myc alters the DNA damage-induced G2/M arrest in human mammary epithelial cells." Br J Cancer **89**(8): 1479-1485.

Vartanian, R., J. Masri, J. Martin, C. Cloninger, B. Holmes, N. Artinian, A. Funk, T. Ruegg and J. Gera (2011). "AP-1 regulates cyclin D1 and c-MYC transcription in an AKT-dependent manner in response to mTOR inhibition: role of AIP4/Itch-mediated JUNB degradation." Mol Cancer Res **9**(1): 115-130.

Wang, Q., Y. Zhou, P. Rychahou, J. W. Harris, Y. Y. Zaytseva, J. Liu, C. Wang, H. L. Weiss, C. Liu, E. Y. Lee and B. M. Evers (2018). "Deptor Is a Novel Target of Wnt/beta-Catenin/c-Myc and Contributes to Colorectal Cancer Cell Growth." Cancer Res **78**(12): 3163-3175.

Yang, Y., K. Xue, Z. Li, W. Zheng, W. Dong, J. Song, S. Sun, T. Ma and W. Li (2018). "c-Myc regulates the CDK1/cyclin B1 dependentG2/M cell cycle progression by histone H4 acetylation in Raji cells." Int J Mol Med **41**(6): 3366-3378.

Ye, F., Y. Chen, T. Hoang, R. L. Montgomery, X. H. Zhao, H. Bu, T. Hu, M. M. Taketo, J. H. van Es, H. Clevers, J. Hsieh, R. Bassel-Duby, E. N. Olson and Q. R. Lu (2009). "HDAC1 and HDAC2 regulate oligodendrocyte differentiation by disrupting the beta-catenin-TCF interaction." Nat Neurosci **12**(7): 829-838.

Zhang, C., M. Lin, R. Wu, X. Wang, B. Yang, A. J. Levine, W. Hu and Z. Feng (2011). "Parkin, a p53 target gene, mediates the role of p53 in glucose metabolism and the Warburg effect." Proc Natl Acad Sci U S A **108**(39): 16259-16264.
